# BioGraphX-RNA: A Universal Physicochemical Graph Encoding for Interpretable RNA Subcellular Localization Prediction

**DOI:** 10.64898/2026.02.23.707573

**Authors:** Abubakar Saeed, Waseem Abbas

## Abstract

RNA subcellular localization is a critical determinant of cellular function. However, current computational approaches often operate as “black boxes,” overlooking the complex interplay among sequence, structure, and physicochemical interactions that govern RNA localization. Building upon the BioGraphX framework originally developed for proteins, we introduce BioGraphX-RNA, a universal physico-chemical graph-encoding framework that provides a structure-informed encoding by translating primary nucleotide sequences into multi-scale interaction graphs using explicit biophysical rules. When combined with frozen RiNALMo embeddings via an interpretable gated fusion layer, BioGraphX-RNA achieves competitive performance with DeepLocRNA and uniquely quantifies the relative contribution of sequence versus structure for each RNA. On human datasets, the gated fusion model attains macro-AUROC values of 0.7575 ± 0.0054 (mRNA), 0.9228 ± 0.0137 (miRNA), and 0.5600 ± 0.0191 (lncRNA). For miRNA, the graph-only model alone reaches 0.9396 ± 0.0045, out-performing both the RiNALMo language model and a RNAfold partition-function graph (0.9139 ± 0.0138), validating the structure-informed proxy hypothesis. In a blind cross-species prediction task on mouse data, the model shows limited zero-shot transfer, indicating that biophysical graph features do not improve cross-species generalization. Gating analysis reveals RNA-type-specific modality reliance, with miRNA exhibiting a near-equilibrium balance between sequence and structure. SHAP-based interpretation suggests potential correlates such as patterned GC content for nuclear retention and structural accessibility for exosome targeting. These advances are achieved with only 2.05 million trainable parameters, aligning with Green AI principles. BioGraphX-RNA demonstrates that explicitly integrating biophysical constraints into graph-based encodings enables accurate and interpretable predictions for structured RNAs, advancing structure-aware RNA biology and laying a foundation for precision medicine.

## 1 Introduction

RNA is the most important part of our understanding of cellular processes, which is significant in the regulation of gene expression, catalysis of biochemical reactions, and translation of genetic information. RNA is largely categorized into three types: messenger RNA (mRNA), long non-coding RNA (lncRNA), and microRNA (miRNA) (Vihinen, 2021; Guo et al., 2015). A major determinant of the biological role of these RNA molecules is their subcellular localization. An example is that lncRNAs are mostly linked to chromatin, and they regulate the transcription and organization of the chromatin. Conversely, mRNAs are found throughout the nucleus and cytoplasm, aiding the movement of genetic information in the form of protein synthesis. These localization mechanisms are governed by complex interactions between cis-regulatory “zip codes” and trans-acting RNA-binding proteins (RBPs). Disruption of RNA localization has also been linked with various disease phenotypes, including cancer (Leucci et al., 2016; Neelamraju et al., 2018; Panda et al., 2018; Zhang et al., 2020), developmental disorders (Nousiainen et al., 2008; Batista et al., 2011; Jao et al., 2017; Okamura et al., 2019), and neuromuscular or neuronal dysfunction (Bassell and Warren, 2008; Dictenberg et al., 2008; Ivy et al., 2010; Baleriola et al., 2014; Didiot et al., 2018).

Traditionally, determining the subcellular location of RNA relied on direct experimental techniques such as fluorescence in situ hybridization (RNA-FISH) and cellular fractionation (Bauman et al., 1980). These approaches remain the gold standards; but they are often expensive, time-consuming, and labor-intensive. To overcome these limitations, the field has increasingly shifted toward computational methods.

Over the past few years, the study of RNA subcellular localization has increasingly shifted toward in silico approaches. This change has been driven by the growing availability of large, well-curated datasets such as RNALocate v3.0 (Wu et al., 2025) and the recent development of RNA foundation models, including RNA-FM (Wang et al., 2024b) and RiNALMo (Penić et al., 2025). Early computational methods primarily relied on manually crafted sequence features combined with conventional machine-learning classifiers. However, more recent approaches have embraced deep neural networks, which can automatically identify patterns associated with RNA localization. For example, DeepLocRNA (Wang et al., 2024a) draws on RNA–protein interaction data to detect localization signals mediated by RNA-binding proteins, while RNALoc-LM (Zeng et al., 2025) leverages pretrained RNA language models to generate embeddings that capture long-range sequence dependencies. Building on these \advances, hybrid models such as LGLoc (Akbari Rokn Abadi et al., 2025), integrate language-model–derived sequence representations with graph neural networks (GNNs) constructed from predicted RNA secondary structures, improving predictions in compartments where structural constraints are particularly important.

Despite these advances, a key limitation remains. Many existing methods operate largely as “black boxes,” relying on phylogenetic patterns or dataset-specific statistical correlations rather than on universal biophysical principles. Consequently, their predictive performance often drops considerably when applied to sequences with low homology or to out-of-distribution data. In addition, a large number of current frameworks treat RNA merely as a linear sequence, overlooking the complex interplay between sequence, structure, and physicochemical interactions that ultimately determines RNA localization.

In this work, we introduce BioGraphX-RNA, a new encoding framework that approximates structural information by converting primary nucleotide sequences into multi-scale interaction graphs using biophysically inspired rules. Building on the structure informed proxy paradigm first developed for the eukaryotic proteome, BioGraphX-RNA extends the BioGraphX (Saeed and Abbas, 2026) architecture to the transcriptomic domain. RNA interaction graphs are constructed directly from sequence data using deterministic physicochemical rules, such as Watson–Crick base pairing and *π*–*π* stacking interactions, eliminating the need for costly and time-consuming three-dimensional structure determination, as shown in Figure 1. Following the principles of Green AI, BioGraphX-RNA outperforms state-of-the-art models like DeepLocRNA (Wang et al., 2024a) while keeping its parameter footprint minimal through a knowledge-integrated design. By incorporating interpretable biophysical features, the framework provides a solid foundation for RNA-focused precision medicine.

**Figure 1:**
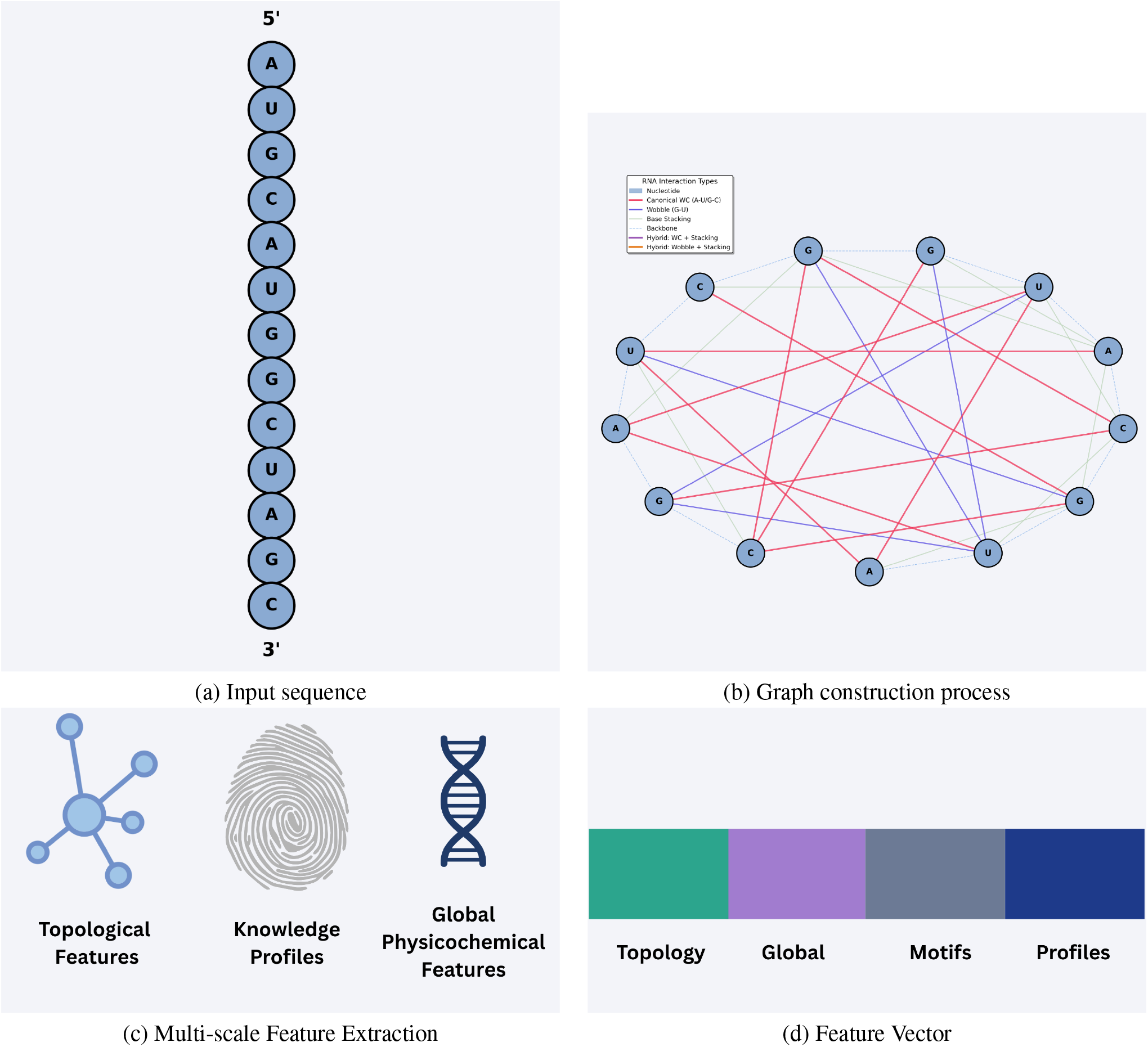
Detailed visualization of BioGraphX-RNA framework. (a) Input RNA sequences for encoding. (b) Construction of rna residue interaction graphs with edges representing biochemical relationships. (c) Multi-scale feature extraction integrating residue-level, global-level, and knowledge-profile features. (d) Feature vector representation ready for downstream classification and explainable analysis.

This work addresses three key questions: (1) whether biophysically grounded graph encodings can serve as a structure-informed proxy without relying on experimental 3D coordinates; (2) whether such representations complement RLMs, and whether any benefit extends across species; and (3) whether explicit biochemical constraints, such as base pairing and *π* −*π* stacking, can yield interpretable predictions that reveal mechanistic insights into RNA trafficking and stability. To this end, we introduce a framework that constructs multi-scale RNA interaction graphs directly from primary sequences using biochemical constraint rules. Unlike existing hybrid methods, which add physical descriptors to pretrained embeddings post hoc, BioGraphX-RNA incorporates biophysical knowledge natively at the encoding stage. The resulting BioGraphX-RNA-Net combines these structural encodings with RiNALMo embeddings through an interpretable gating mechanism, allowing us to quantify the relative contributions of evolutionary signals and biophysical constraints. By keeping the foundation model backbone frozen and training only task-specific parameters, this approach dramatically reduces the number of trainable parameters while maintaining high predictive accuracy, providing a knowledge-driven realization of Green AI for transcriptome-scale prediction.

### 2 Methods

### 2.1 Datasets and Preprocessing

Our research made use of a broad dataset from DeepLocRNA (Wang et al., 2024a), including sequences from nine subcellular locations: Nucleus (13,352), Exosome (22,335), Cytosol (2,587), Cytoplasm (10,026), Ribosome (5,226), Membrane (3,356), ER (1,977), Microvesicle (1,958), and Mitochondrion (33). To ensure a standardized input for deep learning and to identify the most important regulatory signals, mRNA sequences, which are typically much longer, were truncated by retaining only the first 2,500 and last 2,500 nucleotides. In contrast, lncRNA and miRNA sequences were kept in full length. Additionally, all thymine (T) residues were converted to uracil (U) to reflect the biochemical environment of RNA transcripts. The carefully filtered data was then divided into 5-fold subsets according to the distribution of RNA types and sub-cellular locations, for example, genes with multi-labels such as “111000000” (Nucleus, Exosome, and Cytosol) were divided in order to maintain consistency across folds, and an independent benchmarking dataset wasalso set aside to provide an unbiased assessment. This data preprocessing step, including the truncation and T-to-U conversion, was done in the same way for both human and mouse datasets to ensure consistency in methodology across species.

To prevent potential cross-fold homology, we applied CD-HIT-EST Fu et al. (2012) at 80% sequence identity to the combined sequences from all folds. Clusters containing sequences from multiple folds were reassigned entirely to the majority fold. This corrected 82 clusters for miRNA, 6 for lncRNA, and 1 for mRNA. All experiments were repeated on these homology-free splits; results are reported in Supplementary Table 7 and remained highly consistent with the original findings (maximum absolute AUROC change 0.0100, maximum absolute AUPRC change 0.0080), confirming that the CD-HIT-EST filtering had negligible impact on model performance.

To provide a direct comparison with thermodynamically derived structural representations, we implemented two RNAfold-based graph (Lorenz et al., 2011) baselines. For each miRNA sequence, the minimum free-energy (MFE) secondary structure was predicted using RNAfold (ViennaRNA package), and an undirected graph was constructed with nucleotides as nodes and edges representing base pairs from the dot-bracket notation (weight 1.0) and backbone connections between adjacent nucleotides (weight 0.3). A second graph was built from base-pair probabilities obtained via the partition function (RNAfold -p, probability threshold 0.01), where edge weights equal the pairing probabilities. From both graphs, the identical 149 topological, hybrid, knowledge-guided, global biophysical, and frustration features were extracted as in BioGraphX-RNA, and the same MLP classifier and cross-validation splits were used for evaluation.

### 2.2 Overview of BioGraphX-RNA Net Architecture

The BioGraphX-RNA Net architecture consists of three stages. The first part of the architecture, BioGraphX-RNA encoding, is the process of creating biochemically calibrated constraint graphs from RNA sequences to enable the extraction of multi-scale topological, knowledge-guided, and biophysical features. This process uses an adaptive processing approach, smart truncation, and sliding windows to capture the essential 5’ and 3’ terminal regulatory signals and the internal structural motifs. Next, the RiNALMo (Penić et al., 2025) sequence embedding approach is used to extract high-dimensional structural and functional context using pretrained masked language models. The final step of the architecture is the interpretable gated fusion approach, which combines graph-derived biophysical features and deep contextual embeddings using learned gating weights before multi-label sub-cellular localization classification. A schematic representation of this architecture is shown in Figure 2.

**Figure 2:**
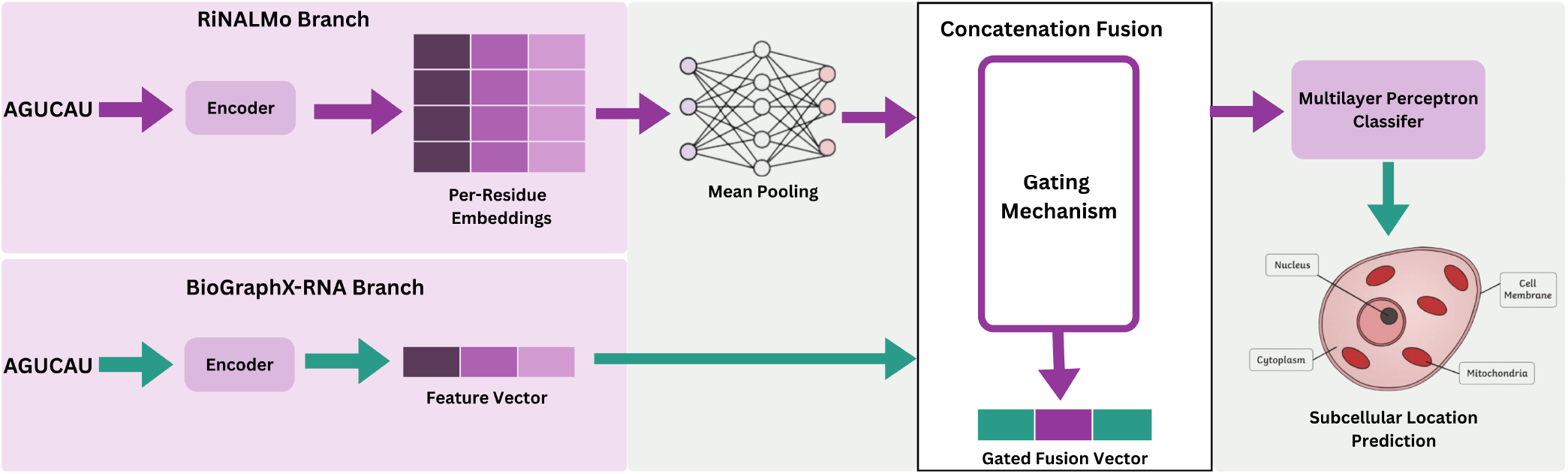
Schematic representation of the BioGraphX-RNA Net computational pipeline. The diagram shows the integration of RiNALMo sequence embeddings and BioGraphX-RNA physics-based features through a central gating mechanism, producing the combined Gated Fusion Vector for downstream classification.

### 2.3 RNA Sequence Embedding and Feature Extraction

The current study employs the RiNALMo model, a state-of-the-art RNA language model, to obtain high-dimensional embeddings, which serve as a dense numerical context for each transcript. Due to the large size of lncRNAs, we adopted an optimized sliding window approach to process variable-length sequences while preserving structural integrity. The sequences were divided into overlapping windows of 2,046 nucleotides with a stride of 1,536 nucleotides (75% overlap). To promote computational efficiency, processing was limited to the first 5,000 nucleotides, prioritizing the most informative regulatory and terminal regions.

To obtain a fixed-length RNA representation from the segment-level embeddings, a mean-pooling approach was adopted. This method helps to guarantee that the structural and functional information present in the entire transcript is retained. The raw embeddings *E*_win_ ∈ℝ ^*L×*1280^ for each window are obtained from the final hidden state of the [CLS] token and are combined into a weighted sequence representation *E*_vec_ through the following operations:

#### Algorithm 1

RNA-BioGraphX Encoding Pipeline

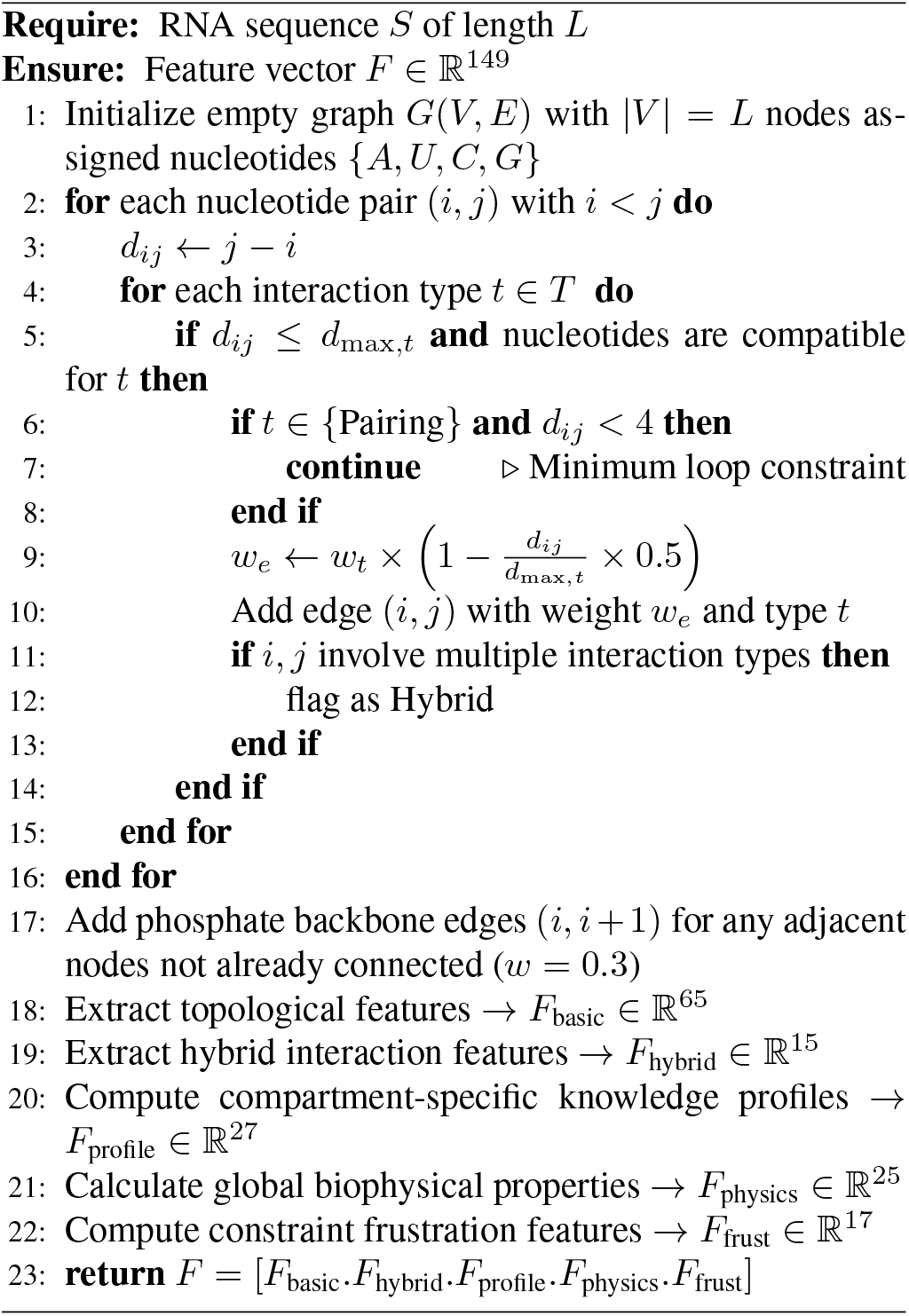

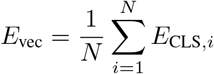

In this representation, *N* is the total number of windows, and *E*_CLS,*i*_ is the class token embedding assigned to the i-th window.

### 2.4 BioGraphX-RNA Encoding Framework

For facilitating transcriptome-scale analysis, the encoding pipeline is equipped with high-performance parallel batch processing capabilities using Joblib with the loky backend, which enhances computational efficiency.

#### 2.4.1 Foundational Biophysical Principles

The BioGraphX -RNA encoding framework translates RNA sequences into multi-scale interaction graphs by applying biochemically calibrated constraint rules. Although language models such as RiNALMo learn statistical patterns from large-scale sequence data, the BioGraphX-RNA framework leverages explicit physical laws that are validated in RNA structural biology. This guarantees the extraction of deterministic and structure-informed features that reflect actual biophysical constraints on RNA folding and localization. The model was particularly adapted from the protein-centric BioGraphX (Saeed and Abbas, 2026) to suit the specific chemical environment of RNA. While the original model used amino acid interactions, the graph model uses nucleotides (A, U, C, G) as nodes and interaction strengths as edges.

#### 2.4.2 Graph Construction Algorithm

For an RNA sequence *S* of length *L*, we build an undirected weighted graph *G*(*V, E*), where vertices *V* = {*v*_1_, *v*_2_, … , *v*_*L*_} correspond to nucleotides, and edges *E* correspond to particular biochemical interactions. To make the computation tractable even for large transcripts, the RNA sequences are first processed using the adaptive truncation approach. The full construction algorithm is presented in Algorithm 1.

#### 2.4.3 Biochemical Interaction Rules

The determination of the particular types of RNA interactions and the setting of the parameters for these interactions were grounded on existing literature in structural biology, such as the Turner rules for RNA structure prediction (Mathews et al., 2004). It must be noted that all the distance thresholds mentioned in this framework are linear sequence separation and not three-dimensional proximity, which enables the model to serve as a proxy based on sequence information only, as shown in Table 1.

**Table 1:**
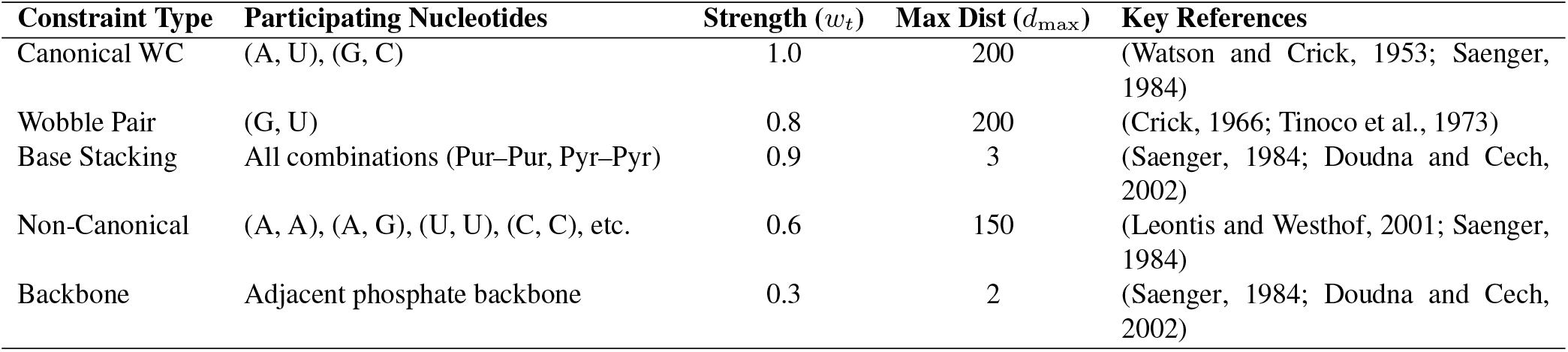
Biochemical interaction types in BioGraphX-RNA.

As illustrated in our framework, these rules encompass the major forces that contribute to RNA stability, from the major stabilizing forces of canonical Watson-Crick base pairing to the particular stabilizing motifs of wobble G-U base pairs and base stacking. The distance thresholds (*d*_max,*t*_) and the strength weights (*w*_*t*_) allow the modeling of the hierarchy of RNA folding and transcript stability. This is done by using the distance thresholds to model the maximum sequence distance over which these forces are active, such as long-range secondary structure interactions, and the weights to model the strength of each type of interaction to the overall biochemical stability of the RNA.

To further model this structural representation, a weight *w*_*t*_ ∈[0, 1] was assigned to each type of interaction and a distance decay function was defined to model the decay of interaction strength with increasing linear sequence distance. The weight *w*_*e*_ of a given edge *E* is given by:

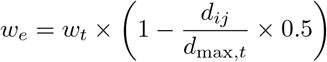

In this model, *d*_*ij*_ is the sequence distance between two nucleotides *i* and *j*, and *d*_max,*t*_ is the maximum allowed distance for a given interaction type *t*. By using these deterministic rules, BioGraphX-RNA models a representation that is grounded in biophysical principles, capturing the competitive process of RNA folding, such as the trade-off between local stacking and long-range pairing, without the need for experimental 3D coordinates. The transition to a 0.5 coefficient is due to the entropic cost of base pairing over longer distances and the prevalence of highly localized structural “zip-code” motifs in RNA. This will ensure that the graph favors stable local secondary structures and, in effect, penalizes distant, less likely conformational interactions. It is important to emphasize that this encoding represents a weighted superposition of biophysical pairing propensities under distance-decay constraints, not a committed secondary or tertiary structure. A single nucleotide may participate in multiple mutually exclusive interaction types simultaneously, reflecting the competitive landscape of RNA folding rather than any single realized conformation.

#### 2.4.4 Hybrid Interaction

One of the major strengths of the BioGraphX-RNA framework is its ability to identify hybrid interactions, where more than one type of interaction, such as base pairing and stacking, occur in close proximity between the same nucleotide pair (Table 2). These interactions are used as high-fidelity signals of specialized structural motifs, which represent regions of high conformational stability. The set of potential hybrid interactions *H* is defined as follows:

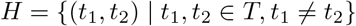

**Table 2:** Hybrid interaction types.

| Hybrid Interaction | Biological Significance |
| --- | --- |
| WC–Stacking | Stabilizes the primary helical stems; core of RNA secondary structure. |
| Wobble–Stacking | Enhances the stability of non-canonical motifs often found in internal loops. |
| Stacking–Backbone | Critical for the formation of tight turns, pseudoknots, and tertiary folds. |
| Non-Canonical–Stacking | Found in specialized regulatory elements and G-quadruplex architectures. |
| Multi-Pairing Hybrid | Occurs at junction points where multiple distal regions converge structurally. |

In the RNA pipeline, we focus on three major hybrid types: Watson-Crick Stacking, Wobble Stacking, and Stacking-Backbone hybrids. When a hybrid interaction is identified, the framework assigns a weight multiplier to the edge, which represents the biochemical stability of these structural “anchors”.

#### 2.4.5 Multi-Scale Feature Extraction

The BioGraphX-RNA engine produces a high-dimensional representation with 149 features that are divided into five different categories: topological, hybrid, knowledge-guided, global biophysical, and constraint frustration (Table 3). This multi-scale representation guarantees that both the macro-scale transcript structure and the micro-scale nucleotide interactions are modeled for sub-cellular localization prediction.

**Table 3:** Feature composition of BioGraphX-RNA encoding framework.

| Feature Scale | # Features | Description | Key Metrics |
| --- | --- | --- | --- |
| Topological | 65 | Network metrics quantifying global and local graph architecture. | Degree statistics, betweenness centrality, path length, modularity, community count. |
| Hybrid | 15 | Co-occurrence patterns of stabilizing interaction types. | WC-Stacking density, regional hybrid clusters, hybrid energy stabilization. |
| Knowledge-Guided | 27 | Profiles derived from known compartment-specific signals. | Motif-based scores for 9 compartments, hybrid compatibility, GC alignment. |
| Global Biophysical | 25 | Whole-transcript biophysical and compositional properties. | GC/AU skew, Shannon entropy, MFE per nucleotide, pairing potential. |
| Constraint Frustration | 17 | Metrics quantifying structural conflicts and local energy variances. | Frustration hotspot density, entropy of frustration, 5' vs 3' structural contrast. |
| **Total** | **149** | Multi-scale RNA representation. | Integrated sequence-structure- features. |

The complete list of 149 features, their mathematical definitions, and logic of feature extraction is provided in Supplementary Material S1.

#### 2.4.6 Sequence Adaptive Processing

To address the high variability in the sizes of the RNA transcripts, ranging from small regulatory RNAs to very large long non-coding RNAs (lncRNAs), we adopted a three-tier adaptive processing approach. This approach is designed to ensure computational scalability while maintaining the essential structural “zipcodes” and terminal signals required for sub-cellular localization, as shown in Supplementary figure Sequences are divided based on their size (*L*) as follows:

- **Short Sequences (***L* ≤1, 000 **nt):** These sequences are processed using their full-length sequences. Due to their relatively small size, the graph building and feature extraction steps are carried out without any loss of information, capturing the entire structural landscape of the RNA.
- **Medium Sequences (**1, 000 *< L* ≤5, 000 **nt):** These sequences are subjected to tri-segmented smart truncation to a fixed length of 1,000 nucleotides. To retain the most biologically important features, we retain the first 30% (5’ terminus), the middle 40% (internal structural features), and the last 30% (3’ terminus). The basis for this split is that the 5’ and 3’ ends are known to contain essential localization signals and protein-binding sites, respectively, while the central regions are known to be rich in structural elements.
- **Long Sequences (***L >* 5, 000 **nt):** For long transcripts, a sliding window strategy is adopted with a window size of 500 nucleotides and a step size of 250 nucleotides. The aggregation of features over the windows is done through an information content-weighted strategy. This strategy weighs motif density and Shannon entropy to focus on “functionally dense” regions, thereby avoiding the dilution of localized regulatory “zipcodes” by unstructured regions of the transcript.

Dynamically adjusting the processing depth according to the length of the transcripts, the BioGraphX-RNA framework preserves a high signal-to-noise ratio over the entire human and mouse transcriptomes.

### 2.5 Hybrid Fusion Architecture

#### 2.5.1 Dual-Branch Network Design

The fusion stage integrates the two complementary representations: the BioGraphX-RNA features and the RiNALMo embeddings. We follow the interpretable gated fusion mechanism as proposed in BioGraphX-Net for proteins (Saeed and Abbas, 2026) , but make a number of simplifications to suit the RNA data. Before fusion, both the embedding vectors are transformed into a common latent space of dimension d=512 to guarantee equal contributions from both branches

##### RNA embedding branch

The mean-pooled RiNALMo embedding *E*_seq_ ∈ℝ^1280^ is transformed using a linear bot-tleneck:

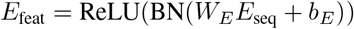

where *W*_*E*_ ≤ ℝ^512*×*1280^ and BN is batch normalization.

##### BioGraphX RNA branch

The 149-dimensional feature vector B is transformed in a single linear layer (in contrast to the three-layer MLP in the protein version):

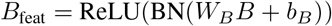

with *W*_*B*_ ∈ℝ^512*×*149^. Preliminary experiments showed that adding additional non-linear layers did not result in any improvement in RNA localization accuracy.

#### 2.5.2 Gated Fusion Mechanism

We keep the gating mechanism as in BioGraphX-Net because of its interpretability and dynamic weighting for each per RNA. The transformed features are then concatenated as follows:

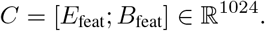

A two-layer gating network computes normalised weights:

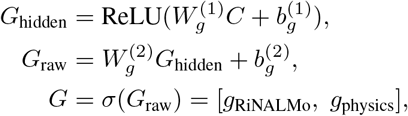

where *σ* is the sigmoid function applied element-wise, and the two gates represent the embedding and physics tasks respectively. The gated feature vector is then computed as follows:

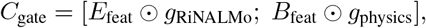

with ⊔ denoting element-wise multiplication.

We note that the gated fusion does not yield a consistent accuracy advantage over simple concatenation (Supplementary Table 8), its primary value lies in enabling per-sample quantification of modality reliance, which is not possible with black-box fusion methods.

#### 2.5.3 Multi-layer Classifier

The last level of classification is a multi-layer perceptron (MLP) that projects the gated features to the subcellular compartment labels. The resulting model has roughly 2.05 million parameters, which is computationally efficient.

#### 2.5.4 Training Protocol and Optimization

The training procedure included the following steps: (1) five-fold cross-validation; (2) saving model checkpoints for each fold based on the maximum validation Macro AUROC; (3) implementing early stopping with a patience of 15 epochs, monitoring validation Macro AUROC; (4) using a batch size of 128; (5) setting the maximum number of training epochs to 100; and (6) applying gradient clipping with a norm of 1.0 after each backward pass. Per-class classification thresholds were optimized by maximizing the Matthews Correlation Coefficient (MCC) for each individual class using a grid search of 50 evenly spaced thresholds from 0 to 1 on each validation fold, following the protocol of DeepLocRNA. All metrics reported in the main tables (Tables 4–6) are computed per-fold using each fold’s own optimized thresholds and are presented as mean ± standard deviation across the five test folds, providing a direct single-model comparison with DeepLocRNA.

**Table 4:** Performance comparison on human mRNA. Best value in each row is highlighted in bold.

| Compartment | BioGraphX-RNA (Ours) |  |  |  | DeepLocRNA* |  |  |  |
| --- | --- | --- | --- | --- | --- | --- | --- | --- |
|  | F1 | MCC | AUROC | AUPRC | F1 | MCC | AUROC | AUPRC |
| Nucleus | $0.7896 \pm 0.0252$ | **$0.4036 \pm 0.0099$** | **$0.7902 \pm 0.0022$** | **$0.8856 \pm 0.0025$** | **0.8341** | 0.3516 | 0.7647 | 0.8681 |
| Exosome | $0.9090 \pm 0.0709$ | **$0.0766 \pm 0.0148$** | **$0.7974 \pm 0.0312$** | **$0.9982 \pm 0.0003$** | **0.9961** | 0.0000 | 0.7537 | 0.9968 |
| Cytosol | **$0.3604 \pm 0.0083$** | **$0.2477 \pm 0.0084$** | $0.7186 \pm 0.0036$ | $0.2946 \pm 0.0108$ | 0.2016 | 0.1627 | **0.7462** | **0.3383** |
| Ribosome | **$0.5735 \pm 0.0083$** | **$0.3572 \pm 0.0104$** | **$0.7621 \pm 0.0038$** | **$0.5699 \pm 0.0034$** | 0.3860 | 0.2460 | 0.7564 | 0.5509 |
| Membrane | **$0.4478 \pm 0.0252$** | **$0.3251 \pm 0.0061$** | **$0.7592 \pm 0.0040$** | **$0.4667 \pm 0.0062$** | 0.3001 | 0.2569 | 0.7532 | 0.4337 |
| ER | **$0.3336 \pm 0.0137$** | **$0.2418 \pm 0.0164$** | $0.7174 \pm 0.0032$ | **$0.3070 \pm 0.0131$** | 0.0702 | 0.0474 | **0.7217** | 0.2836 |
| Macro Avg$^{\ddagger}$ | **$0.5690 \pm 0.0146$** | **$0.2753 \pm 0.0041$** | **$0.7575 \pm 0.0054$** | **$0.5870 \pm 0.0017$** | 0.4647 | 0.1774 | 0.7493 | 0.5786 |

\* DeepLocRNA values from Supplementary Table 1.
$^{\ddagger}$ Macro averages for DeepLocRNA computed over the six reported compartments.

### 2.6 Explainability Framework

To interpret BioGraphX-RNA Net predictions, we employed the same multi-pronged explainability framework described in our BioGraphX protein work, comprising gate analysis, SHAP feature importance, and knowledge-guided profiling.

### 2.7 Evaluation Metrics

We evaluated all models using four metrics: F1 score, Matthews Correlation Coefficient (MCC), Area Under the Receiver Operating Characteristic curve (AUROC), and Area Under the Precision-Recall curve (AUPRC). Metrics are reported as mean ± standard deviation across the five cross-validation test folds. AUROC and AUPRC are threshold-independent; F1 and MCC are computed using per-class decision thresholds optimized by MCC maximization on each validation fold, following the protocol of DeepLocRNA.

### 2.8 Computational efficiency (Green AI)

All experiments were conducted on the Kaggle platform using a single kaggle NVIDIA T4 GPU (16 GB). RiNALMo embeddings were generated once for the full human dataset and required approximately 7.8 hours. Mouse embeddings were generated separately with a similar setup. The resulting embeddings and 149 BioGraphX graph features were stored and reused across all downstream experiments. Classifier training (2.05 million trainable parameters) completed in less than 5 minutes per five-fold cross-validation. Total end-to-end compute for a full reproduction is approximately 10 GPU-hours. Because the language model backbone remains frozen and graph features are derived from lightweight deterministic rules, the marginal cost of additional experiments is negligible, aligning the framework with Green AI principles.

## 3 Results

We assess BioGraphX-RNA on the DeepLocRNA benchmark, showing improvements across all types of RNAs and cellular compartments. Moreover, we set the first blind cross-species benchmark by evaluating human-trained models on fully held-out mouse data, a stricter test than before, where the target species’ sequences were inadvertently included in the training set. To provide context, we refer to the performance of DeepLocRNA as reported in their supplementary materials. For mRNA, we report the performance of the “instructive fine-tuning + weighting” model (Supplementary Table 1 in (Wang et al., 2024a)); for miRNA, we report the performance of the unified model on the miRNA subset (Supplementary Table 2); and for lncRNA, we report the performance of their specialized lncRNA model (Supplementary Table 6). Notably, the mouse performance of DeepLocRNA is not directly comparable since mouse sequences were included in their training set; we address cross-species generalization separately.

### 3.1 Ablation Study

We dissected the contribution of each component by comparing four configurations, reporting per-fold mean ± standard deviation across the five cross-validation folds (Supplementary Table S8): (i) *RiNALMo-only*, frozen foundation model embeddings passed to the classifier; (ii) *graph-only*, our 149-dimensional biophysical graph features alone, using the same MLP classifier; (iii) *simple concatenation*, concatenation of graph features and RiNALMo embeddings; and (iv) *full gated fusion*, our proposed model with interpretable gating.

#### Graph-only as a structure-informed lightweight encoder

The graph-only model, which relies solely on explicit biophysical rules (Watson–Crick pairing, stacking, wobble, backbone, distance decay), achieved the highest single-modality performance for miRNA (macro AUROC 0.9396 ± 0.0045), exceeding both RiNALMo-only (0.9157 ± 0.0069) and the gated fusion (0.9228 ± 0.0137). This demonstrates that the hand-crafted physicochemical features alone capture a substantial fraction of the localization signal for small, highly structured RNAs. For mRNA and lncRNA, the graph-only model is considerably weaker than the language model embeddings, indicating that the structural signal is less informative for longer, less structurally constrained transcripts. The graph-only representation is fully interpretable: its 149 features are grouped into topological, hybrid, knowledge-guided, global biophysical, and frustration categories (Supplementary Material S1), each directly linked to biochemical principles. The graph-only model is also extremely lightweight (149 features, no large language model), making it suitable for high-throughput screening on standard CPUs.

#### Combining modalities yields modest improvements for mRNA, but not for lncRNA

For mRNA, simple concatenation of graph features and RiNALMo embeddings improved macro AUROC from 0.7572 ± 0.0074 (RiNALMo-only) to 0.7581 ± 0.0028, and macro AUPRC from 0.5910 ± 0.0026 to 0.5879 ± 0.0026. For lncRNA, however, RiNALMo-only alone achieved the highest AUROC (0.5889 ± 0.0247); concatenation with graph features yielded 0.5654 ± 0.0049, and the gated fusion 0.5600 ±0.0191, both slightly below the language-model baseline. This indicates that for lncRNAs, the frozen language model embeddings already capture the available localization signal, and the addition of biophysical graph features does not improve and may slightly degrade performance. For miRNA, graph features alone already surpass the language model; concatenation with RiNALMo embeddings yields 0.9149 ± 0.0152, which remains below the graph-only result, further underscoring the primacy of structural information for this RNA type.

#### Interpretable gating versus simple concatenation

The gated fusion achieved performance comparable to simple concatenation across all RNA types, with differences well within one standard deviation of the fold-level means: for mRNA, ΔAUROC = −0.0006, for miRNA, gated fusion (0.9228 ± 0.0137) slightly exceeded concatenation (0.9149 ± 0.0152), for lncRNA, ΔAUROC = −0.0054. While the gating mechanism did not provide a significant accuracy advantage, it enables the quantification of modality reliance at the individual RNA level. As shown in Section 3.5 and Figure 3, gate values reveal that miRNA depends nearly equally on sequence and structure, whereas mRNA relies more on sequence. This interpretability is unique to the gated architecture and generates testable biological hypotheses. Therefore, even when accuracy is on par with simpler fusion methods, the gated fusion adds a layer of interpretable insight that is absent from black-box concatenation.

**Figure 3:**
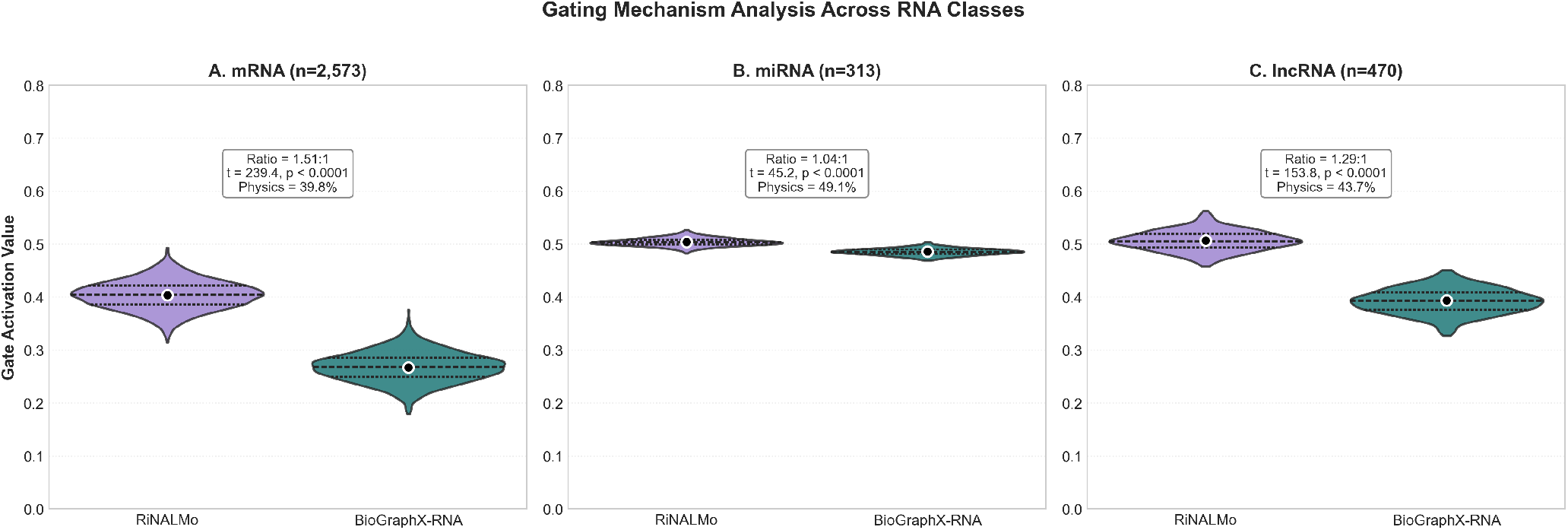
Gating mechanism analysis across RNA classes. Violin plots showing the distribution of gate activation values for RiNALMo embeddings (purple) and BioGraphX-RNA physics features (teal). Black dots indicate means; inner lines show quartiles. **(A)** mRNA (n=2,573): Embedding-dominated with physics contributing 39.8%. **(B)** miRNA (n=313): Near-perfect balance with exceptionally narrow distributions, reflecting absolute structural requirements. **(C)** lncRNA (n=470): Intermediate integration with wider distributions capturing functional diversity.

In summary, the ablation confirms that: (1) the biophysical graph encoding is a powerful standalone representation for structured RNAs, most notably miRNA, where it surpasses both the language model and thermodynamic folding baselines; (2) for mRNA, combining graph features with sequence embeddings offers modest gains, while for lncRNA the language model alone is sufficient; and (3) the gated fusion is preferred when interpretability of modality reliance is desired, while simple concatenation offers a slightly simpler alternative without meaningful accuracy loss.

### 3.2 Structure-Informed Proxy Validation Against RNAfold

A central question for any heuristic graph-encoding method is whether it captures genuinely informative structural signals or merely approximates information already represented by thermodynamic folding methods. To address this directly, we compared the graph-only variant of BioGraphX-RNA against two RNAfold-derived graph baselines using the same MLP classifier and the same cross-validation splits.

We selected miRNA for this comparison because it represents the most favorable setting for folding-based approaches, miRNA precursor sequences are short, and RNAfold predictions are generally most reliable in this regime. If BioGraphX-RNA could not outperform a folding-based graph representation under these conditions, the claim that it acts as an effective structure-informed proxy would be substantially weakened.

Two RNAfold graphs were constructed. The first uses the minimum free-energy (MFE) dot-bracket structure, with binary edges for backbone and base pairs. The second uses base-pair probabilities from the partition function, producing a weighted graph that reflects the full thermodynamic ensemble rather than a single committed structure. From both graphs, the same 149 topological, hybrid, knowledge-guided, global biophysical, and frustration descriptors were extracted as in BioGraphX-RNA, allowing a comparison at identical feature dimensionality.

BioGraphX-RNA graph-only achieved a macro AUROC of 0.9396 ± 0.0045. The MFE graph yielded 0.8957 ± 0.0184, and the partition-function graph reached 0.9139 ± 0.0138. The partition-function representation substantially improved over the MFE graph, confirming that probabilistic ensemble information aids localization prediction. Nevertheless, BioGraphX-RNA retained a ΔAUROC of 0.0257 over the partition-function baseline, a margin observed in all five cross-validation folds.

The remaining gap reflects the richer interaction vocabulary of the BioGraphX graph. RNAfold graphs, whether MFE or partition-function, model a single interaction type (base-pairing). In contrast, BioGraphX-RNA encodes canonical Watson–Crick pairing, wobble interactions, *π*–*π* stacking, non-canonical interactions, and backbone connectivity, each with biophysically calibrated weights and distance-dependent decay (Table 1, Algorithm 1). These multiple interaction types are represented as a weighted superposition that captures local structural propensities beyond what a single base-pair probability can express. Furthermore, the frustration features (Supplementary Material S1) quantify conflicts between competing interaction potentials, information that has no direct analog in a thermodynamic folding graph.

The monotonic progression, MFE (0.8957) →partition function (0.9139) →BioGraphX-RNA (0.9396), demonstrates that each additional layer of local conformational information improves localization prediction.

That BioGraphX-RNA outperforms the partition-function graph, which encodes the full thermodynamic ensemble of base-pairing probabilities, provides strong evidence that the heuristic multi-interaction encoding captures genuine structural signals beyond those available from state-of-the-art folding algorithms. This finding supports the use of BioGraphX-RNA as a structure-informed proxy for RNA localization prediction, particularly for small, highly structured RNAs where the advantage is largest.

### 3.3 Performance on Human RNAs

#### Mrna

The results for mRNA are summarized in Table 4 for the six subcellular compartments. BioGraphX-RNA (gated fusion) achieves a macro-AUROC of 0.7575 ± 0.0054, a modest improvement over DeepLocRNA’s reported macro-AUROC of 0.7493. The largest gains are observed in per-class F1 and MCC, particularly for Cytosol and ER, indicating that the biophysical graph features provide complementary information for compartments where sequence-only signals are weak. All metrics are reported with per-fold standard deviations, which are not available for DeepLocRNA.

#### miRNA

For miRNA, our model was evaluated on five compartments (Table 5). BioGraphX-RNA achieves a macro-AUROC of 0.9228 ± 0.0137, substantially higher than DeepLocRNA’s unified model (0.8681). The improvement is most pronounced in Cytoplasm and Mitochondrion, despite the latter having only 33 training samples. These results demonstrate that biophysically-calibrated structural features are particularly effective for small, highly structured RNAs, where folding constraints are functionally critical.

**Table 5:**
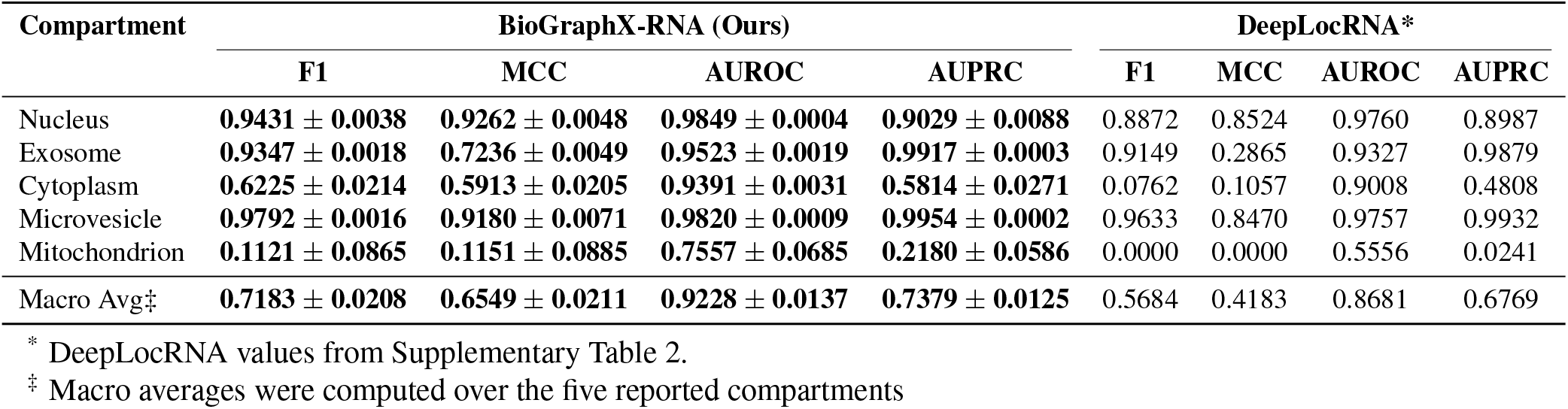
Performance comparison on human miRNA. Best value in each row is highlighted in bold.

| Compartment | BioGraphX-RNA (Ours) |  |  |  | DeepLocRNA* |  |  |  |
| --- | --- | --- | --- | --- | --- | --- | --- | --- |
|  | F1 | MCC | AUROC | AUPRC | F1 | MCC | AUROC | AUPRC |
| Nucleus | **0.9431 ± 0.0038** | **0.9262 ± 0.0048** | **0.9849 ± 0.0004** | **0.9029 ± 0.0088** | 0.8872 | 0.8524 | 0.9760 | 0.8987 |
| Exosome | **0.9347 ± 0.0018** | **0.7236 ± 0.0049** | **0.9523 ± 0.0019** | **0.9917 ± 0.0003** | 0.9149 | 0.2865 | 0.9327 | 0.9879 |
| Cytoplasm | **0.6225 ± 0.0214** | **0.5913 ± 0.0205** | **0.9391 ± 0.0031** | **0.5814 ± 0.0271** | 0.0762 | 0.1057 | 0.9008 | 0.4808 |
| Microvesicle | **0.9792 ± 0.0016** | **0.9180 ± 0.0071** | **0.9820 ± 0.0009** | **0.9954 ± 0.0002** | 0.9633 | 0.8470 | 0.9757 | 0.9932 |
| Mitochondrion | **0.1121 ± 0.0865** | **0.1151 ± 0.0885** | **0.7557 ± 0.0685** | **0.2180 ± 0.0586** | 0.0000 | 0.0000 | 0.5556 | 0.0241 |
| Macro Avg‡ | **0.7183 ± 0.0208** | **0.6549 ± 0.0211** | **0.9228 ± 0.0137** | **0.7379 ± 0.0125** | 0.5684 | 0.4183 | 0.8681 | 0.6769 |

\* DeepLocRNA values from Supplementary Table 2.
‡ Macro averages were computed over the five reported compartments

#### lncRNA

Long non-coding RNAs remain challenging to predict due to their diverse sequences, low conservation, and heterogeneous functions. Following DeepLocRNA, we evaluate on four compartments (Nucleus, Exosome, Cytosol, Membrane; Table 6). BioGraphX-RNA achieves a macro-AUROC of 0.5600 ± 0.0191, slightly lower than DeepLocRNA’s 0.5786, but with substantially higher per-class F1 on Nucleus and Cytosol. The lower macro-AUROC reflects the inherent difficulty of lncRNA localization: the frozen language model alone already yields the best discriminative performance (Supplementary Table S8), and the addition of graph features primarily improves decision-threshold-dependent metrics rather than ranking-based metrics. This pattern suggests that the biophysical features help separate positive from negative examples within individual compartments even when overall ranking ability is dominated by sequence signals.

**Table 6:** Performance comparison on human lncRNA. Best value in each row is highlighted in bold.

| Compartment | BioGraphX-RNA (Ours) |  |  |  | DeepLocRNA* |  |  |
| --- | --- | --- | --- | --- | --- | --- | --- |
|  | F1 | MCC | AUROC | AUPRC | MCC | AUROC | AUPRC |
| Nucleus | **0.4199 ± 0.0062** | **0.1174 ± 0.0177** | **0.6058 ± 0.0351** | **0.3469 ± 0.0298** | 0.0156 | 0.5443 | 0.3027 |
| Exosome | **0.7435 ± 0.1819** | **0.0292 ± 0.0379** | 0.5821 ± 0.0369 | **0.9822 ± 0.0024** | 0.0000 | **0.5832** | 0.9815 |
| Cytosol | **0.1142 ± 0.0483** | **0.0290 ± 0.0522** | 0.5502 ± 0.0327 | 0.0864 ± 0.0208 | 0.0000 | **0.5932** | **0.1004** |
| Membrane | **0.0522 ± 0.0094** | **-0.0037 ± 0.0242** | 0.5018 ± 0.0350 | 0.0405 ± 0.0057 | 0.0000 | **0.5938** | **0.0658** |
| Macro Avg‡ | 0.3324 ± 0.0469 | **0.0430 ± 0.0099** | 0.5600 ± 0.0191 | **0.3640 ± 0.0104** | 0.0039 | **0.5786** | 0.3626 |

\* DeepLocRNA values from Supplementary Table 6.
‡ Macro averages were computed over the four reported compartments.

### 3.4 Blind Cross-Species Transfer

To test whether biophysical graph features capture evolutionarily conserved localization signals, we trained our model exclusively on human data and evaluated it on a completely independent mouse dataset. Unlike DeepLocRNA, which included mouse sequences in its training set, this represents a strict zero-shot cross-species test. We report macro-averaged and per-compartment AUROC and AUPRC.

Table 7 summarizes the results. The macro AUROC for mRNA is 0.4889, near the theoretical chance level of 0.5, yet clear compartment-specific differences are visible. Nuclear localization is the most conserved signal (mRNA Nucleus AUROC 0.6753, AUPRC 0.6531; lncRNA Nucleus AUROC 0.6772, AUPRC 0.6299), indicating that sequence features relevant to nuclear retention are partially shared between human and mouse. In contrast, cytoplasmic mRNA localization is predicted below chance (AUROC 0.2907), suggesting that cytoplasmic targeting signals are highly species-specific or require contextual information not present in the sequence alone.

**Table 7:** Blind mouse evaluation results across different RNA types.

| RNA Type | Compartment | AUROC | AUPRC |
| --- | --- | --- | --- |
| mRNA | Nucleus | 0.6753 | 0.6531 |
|  | Exosome | 0.5006 | 0.1939 |
|  | Cytoplasm | 0.2907 | 0.2501 |
|  | **Macro Avg** | **0.4889** | **0.3657** |
| miRNA | Nucleus | 0.4984 | 0.1991 |
|  | Exosome | 0.5489 | 0.8716 |
|  | Mitochondrion | 0.5113 | 0.2988 |
|  | **Macro Avg** | **0.5195** | **0.4565** |
| lncRNA | Nucleus | 0.6772 | 0.6299 |
|  | Exosome | 0.4095 | 0.4422 |
|  | **Macro Avg** | **0.5434** | **0.5361** |

For miRNA, the overall macro AUROC is 0.5195. Exosome-targeting signals show the strongest cross-species transfer (AUROC 0.5489, AUPRC 0.8716), while nucleus and mitochondrion predictions remain near chance.

For lncRNA, macro AUROC reaches 0.5434, driven primarily by nuclear localization (0.6772). Exosome predictions are below chance (0.4095), reflecting the extreme sequence and functional diversity of lncRNAs and the limited transferability of the learned features.

Across all RNA types, the frozen RiNALMo-only baseline and the graph-only model yield comparable cross-species performance (Supplementary Table 9), with macro AUROC values close to chance (mRNA ≈0.49, miRNA ≈0.51; lncRNA ranges from 0.50 to 0.56 depending on the configuration, with RiNALMo-only performing modestly better than the graph-only model). This confirms that biophysical graph features calibrated on human-specific compositional biases do not improve cross-species generalization. The one compartment where a modest above-chance signal is consistently observed is nuclear localization (mRNA Nucleus AUROC ≈0.66–0.69; lncRNA Nucleus AUROC ≈0.57–0.69), and this signal is present in both the sequence embeddings and the graph features, with neither modality providing a clear advantage. Overall, this blind evaluation demonstrates that cross-species RNA localization prediction remains a challenging open problem, with only nuclear retention signals showing limited transferability.

### 3.5 Explainability Analysis

#### 3.5.1 Gating Analysis

The gating function in BioGraphX-RNA dynamically weights the RiNALMo embeddings and BioGraphX graph features for each RNA molecule. Examination of the gate values on the test sets reveals the balance in reliance on biophysical constraints across different RNA types and cellular compartments.

For mRNA, the average physics gate value is 0.267 (39.8% of total gate weight), with the embedding gate at 0.404 (60.2%), as shown in Figure 3A. Physics contribution shows minimal variation across compartments, ranging from 39.5% in Cytosol to 40.0% in Endoplasmic Reticulum. This consistent 40% physics contribution suggests that, in our model, biophysical constraints act as a complementary validation signal even for mRNAs, where sequence signals are dominant.

For miRNA, the physics gate is substantially higher, averaging 0.486 (49.1%), with the embedding gate at 0.504 (50.9%) Figure 3B. This near-perfect balance is consistent with the well-established structural requirements of miRNA biology, including precursor hairpins for DICER processing and conformational specificity for RISC loading and vesicular sorting (Zhang et al., 2004; Bernstein et al., 2001; Grishok et al., 2001). Compartment-specific analysis reveals the highest physics reliance in Mitochondrion (49.3%) and Microvesicle (49.2%), consistent with specialized structural requirements for trafficking to these organelles. Notably, the exceptionally narrow distribution indicates that structural validation is universally required across all miRNAs, regardless of destination.

For lncRNA, the physics gate averages 0.394 (43.7%), with the embedding gate at 0.507 (56.3%). This intermediate position between mRNA (39.8% physics) and miRNA (49.1% physics) reflects the functional and structural diversity of lncRNAs. Compartment-specific analysis reveals subtle but informative variation: cytoplasmic lncRNAs show the highest physics dependence (44.0%), consistent with scaffold functions requiring specific conformations (Noh et al., 2018), while membrane-associated lncRNAs show the lowest (43.3%). The wider distribution compared to miRNAs captures the heterogeneity of lncRNA structure and function, as illustrated in Figure 3C.

These findings demonstrate that BioGraphX-RNA Net effectively learns to integrate evolutionary information with biophysical constraints, and that the gating mechanism dynamically adjusts to the distinct characteristics of each RNA type and subcellular compartment.

#### 3.5.2 SHAP Analysis

To interpret which biophysical features drive BioGraphX-RNA predictions, we performed SHAP analysis on the RNA test set. Given the high-dimensional feature space (149 features), we normalized SHAP values within each feature to the [−1, 1] range, preserving the direction of contribution (positive = increases probability, negative = decreases probability) while enabling cross-compartment comparison. The SHAP analyses described below identify correlational patterns between specific biophysical features and subcellular compartments based on model predictions. SHAP values quantify the contribution of each feature to the model’s output for a given prediction, but they do not imply causality. These associations are therefore hypothesis-generating rather than direct evidence of biological mechanisms. Experimental validation is required to confirm any proposed functional roles.

#### 3.5.3 mRNA

##### Nucleus is associated with patterned GC distribution

For nuclear mRNA localization, the top features were GC Autocorrelation Lag1 (positive) and Five Prime GC (negative) (Fig. 4). This suggests that the model weights periodic GC spacing more heavily than absolute GC content, a pattern consistent with the known roles of GC-rich structural motifs such as Alu repeats (Lubelsky and Ulitsky, 2018) and SIRLOIN elements (Yao et al., 2021) in nuclear retention.

**Figure 4:**
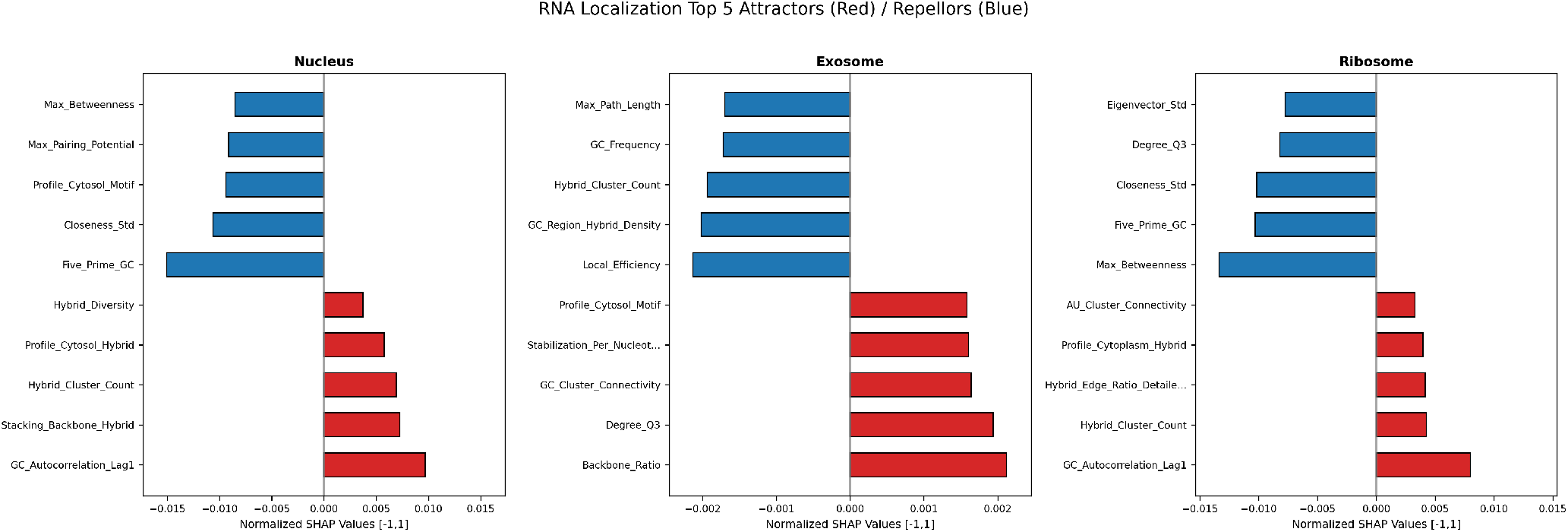
Top five organelle-specific features for mRNA ranked by feature importance values.

##### Exosome targeting correlates with an “anti-structure” profile

Exosome localization was associated with high Backbone Ratio (indicating accessible, unstructured regions) and low Local Efficiency, GC Region Hybrid Density, and Hybrid Cluster Count (indicating fewer protective structural elements). Notably, Profile Exosome Motif, which encodes AU-rich elements (AREs), did not rank among the top features. This pattern suggests that structural accessibility, rather than ARE content alone, is the dominant correlate in our model. The hypothesis that loss of protective structure facilitates exosome targeting is consistent with observations that ARE-mRNAs accumulate in exosome granules upon decay inhibition (Lin et al., 2007), but requires independent experimental validation.

##### Ribosome association correlates with periodic GC patterns and structural heterogeneity

Ribosome-associated mRNAs were positively associated with GC Autocorrelation Lag1 and Hybrid Cluster Count, and negatively associated with Max Betweenness, Closeness Std, and Eigenvector Std. This suggests that periodic GC patterns and hybridized structural elements correlate with ribosomal association in the model, while high topological heterogeneity correlates with exclusion. These associations are consistent with observations of ribosomal structural dynamics (Helena-Bueno et al., 2025; Alejo et al., 2024), but causal links remain to be tested.

#### 3.5.4 miRNA

##### Nucleus is associated with high-degree connectivity and GC-rich hybrid regions

Nuclear-localized miRNAs were positively associated with Degree Q3, Degree Q1, and GC Region Hybrid Density (Fig. 5), indicating that both high and low network connectivity and dense GC-rich hybrid interactions correlate with nuclear retention in the model. Features reflecting structural plasticity, such as Frustration Entropy and Community Size Std, also appeared among the top predictors. These associations are consistent with proposed roles of Argonaute nuclear import (Abaturov and Babych, 2021) and HEN2-mediated regulation (Vigh et al., 2022).

**Figure 5:**
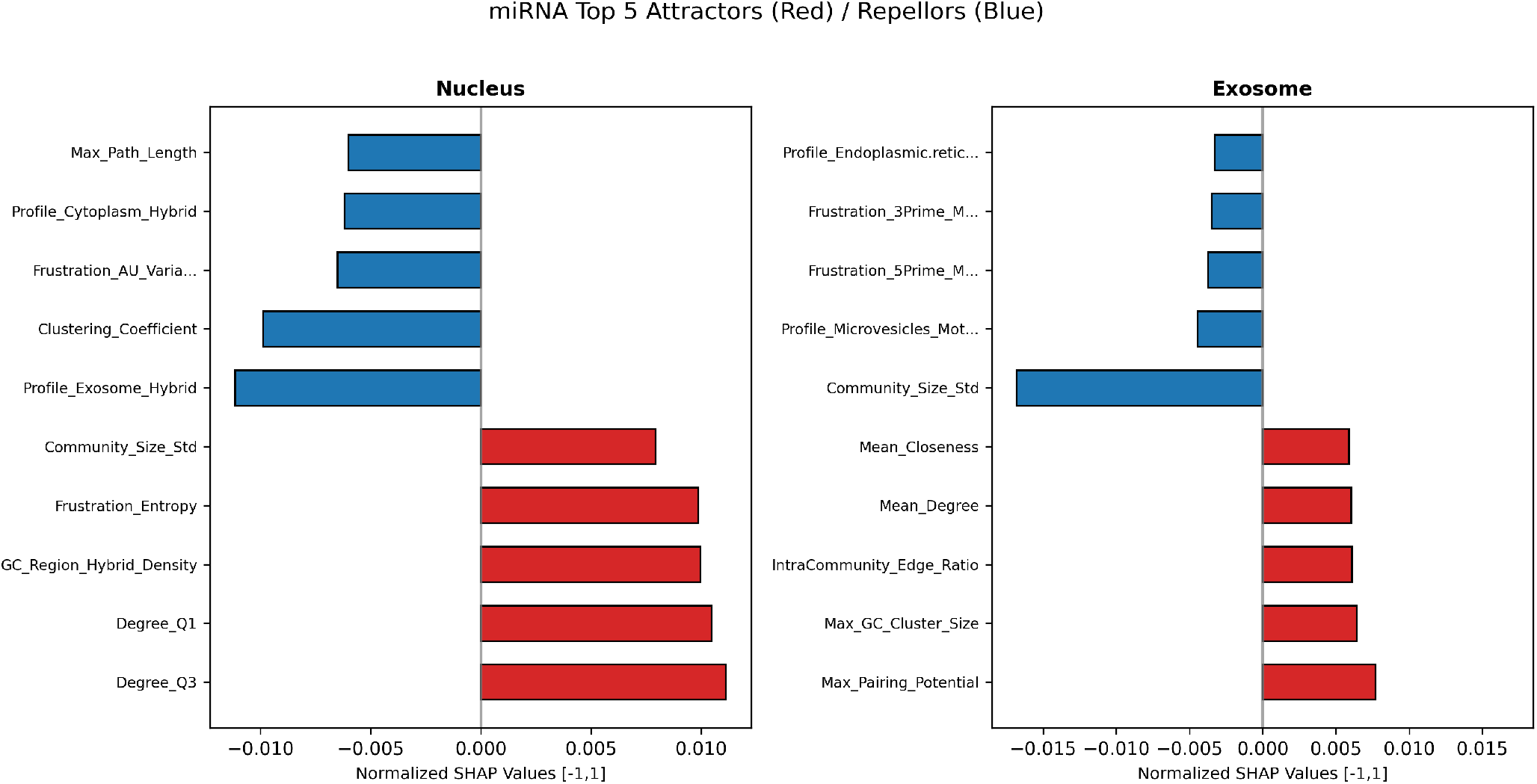
Top five organelle-specific features of miRNA ranked by feature importance values 13.

**Figure 6:**
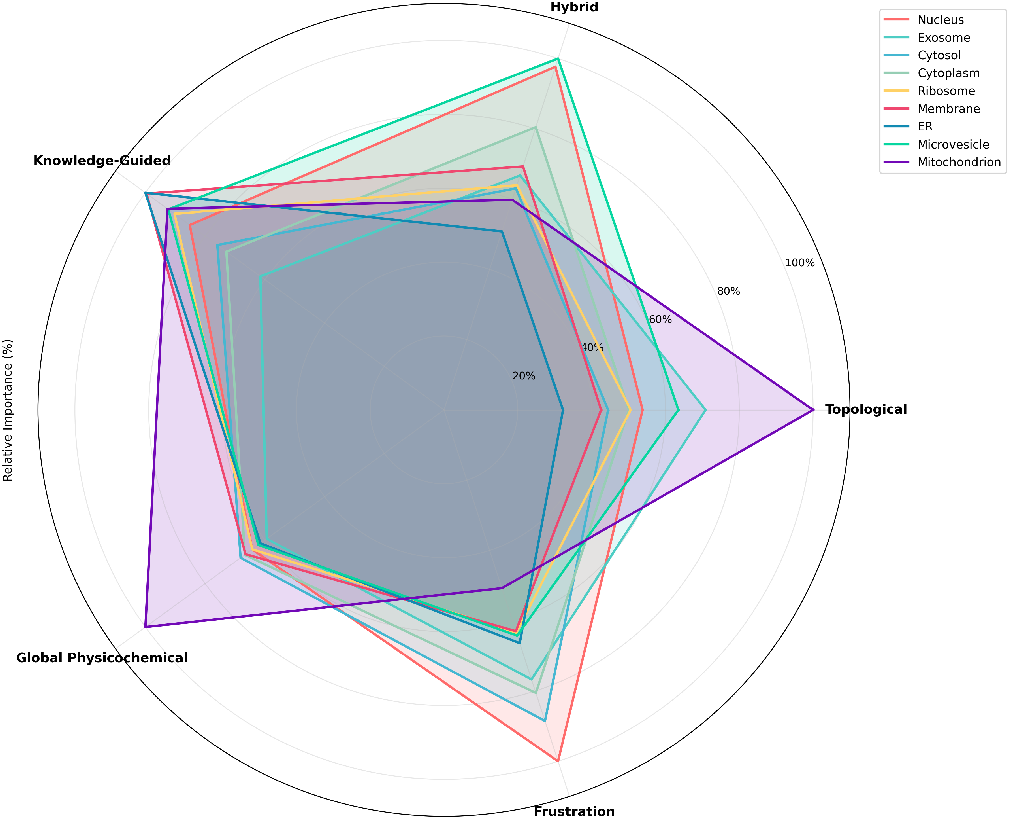
Radar chart summarizing the contribution of five feature categories across all RNA types.

##### Exosome localization correlates with pairing potential and GC cluster architecture

Exosome-localized miR-NAs were associated with Max Pairing Potential, Max GC Cluster Size, and IntraCommunity Edge Ratio, suggesting that high base-pairing propensity, longer GC-rich regions, and internally dense connectivity are preferential correlates of exosome uptake in the model. This pattern is broadly consistent with the binding preferences of exosome-sorting RNA-binding proteins such as hnRNPA2B1 and FUS (Garcia-Martin et al., 2022; Villarroya-Beltri et al., 2013), although the SHAP associations alone do not confirm a direct sorting mechanism.

#### 3.5.5 lncRNA

SHAP analysis of lncRNA predictions revealed generally weak feature contributions (normalized SHAP magnitudes *<* 0.01), consistent with the low overall performance for this RNA type (macro AUROC 0.5600 ± 0.0191). The only compartment with discernible structural correlates was the nucleus, where Std Pairing Potential and Frustration Hotspot-Count appeared among the top features, suggesting that structural variability and localized folding conflicts are weakly associated with nuclear localization.

#### 3.5.6 Feature Category Contributions Across RNA Types

Feature category importance analysis for subcellular compartments indicates specific patterns of biophysical signal usage as a function of cellular target. Topological features, which describe graph architecture in terms of degree distributions, centrality, and community structure, were found to have the highest relative importance in mitochondria, exosomes, and microvesicles, consistent with the importance of interaction networks and structural hotspots in targeting to these specialized organelles. By contrast, topological features had very low importance in the endoplasmic reticulum, suggesting that ER targeting is more sequence motif- and linear signal-based.

Hybrid patterns, representing the co-occurrence of multiple interaction types, were predominantly found in nucleus, microvesicles, and cytoplasm, as expected for these compartments, which need stable structural “anchors” for retention or trafficking.

Knowledge-based features, computed from known compartment-specific sequence motifs, demonstrated a rather similar distribution across compartments, with a slight increase in ribosome, membrane, and ER. The rather small differences indicate that while linear motifs play a role in localization, they do so in combination with structural patterns rather than being the primary determinants.

Global physicochemical properties, such as GC content, dinucleotide composition, and global stability measures, were most important in mitochondria, cytosol , and cytoplasm. The prominent increase in mitochondria reflects the specific needs of mitochondrial import.

Frustration measures, reflecting local structural conflicts and conformational diversity, were most important in nucleus and cytosol, with prominent increases also in exosomes and cytoplasm. The prominent role of frustration in nuclear transcripts is consistent with the model that structural conflict domains are protein interaction platforms for architectural lncRNAs and nuclear retention factors. The high frustration importance in cytosol indicates that conformational flexibility is important for recognition by cytosolic RNA-binding proteins and entry into degradation or translation pathways.

Notably, mitochondria showed the most distinctive set of characteristics: the highest topological and overall physicochemical importance, but lowest frustration. This distinctive set of characteristics is consistent with the specialized needs of mitochondrial RNA import, which requires stable structural elements for membrane passage while avoiding conformational diversity that could hinder uptake into organelles. The near-complementary distribution of characteristics between topological properties and frustration in different compartments, where nucleus and cytosol prefer frustration over topology, mitochondria and exosomes over frustration, demonstrates a deep trade-off between structural heterogeneity (frustration) and structural complexity (topology) in RNA fate decisions.

Taken together, these results show that different subcellular compartments impose distinct biophysical constraints: nucleus and cytosol prefer conformational flexibility and structural heterogeneity (high frustration), whereas mitochondria and exosomes prefer complex interaction networks and structural stability (high topology). This compartment-specific use of characteristics offers a systems-level perspective on how RNA structure encodes information for trafficking.

## 4 Discussion

RNA subcellular localization prediction is still a fundamental problem in computational biology with broad implications for the understanding of gene regulation, cellular function, and disease. In this study, we proposed BioGraphX-RNA, a new approach that provides a structure-informed proxy by transforming primary RNA sequences into multi-scale interaction graphs based on explicit biophysical models. By combining these graph-based features with sequence embeddings from the RiNALMo foundation model using an interpretable gated fusion approach, we show that biophysical models can not only reach state-of-the-art predictive accuracy but also offer mechanistic explanations for the structural basis of RNA localization. Our results answer the three research questions raised in the introduction and thus show the BioGraphX-RNA approach to be a general tool for transcriptome-wide analysis.

### 4.1 Biophysically Grounded Encodings Provide a Structure-Informed Proxy

The first question we addressed was whether biophysically informed graph encodings could provide a useful signal without experimental three-dimensional coordinates. The answer is clearly affirmative. The RNAfold comparison (Section 3.2) demonstrates that BioGraphX-RNA captures structural information beyond thermodynamic folding representations, with the graph-only model outperforming the partition-function baseline (0.9396 vs. 0.9139 macro AUROC). The ablation study (Section 3.1) further shows that these graph features are most informative for miRNA, the most structure-dependent RNA type, where the graph-only model surpasses even the RiNALMo language model. For mRNA and lncRNA, the biophysical features provide complementary information that improves decision-threshold-dependent metrics, while the overall ranking ability remains dominated by sequence embeddings.

Independent cross-domain validation comes from our published BioGraphX protein study (Saeed and Abbas, 2026), where the same encoding principles, applied to a 3D-dependent protein solubility task—achieved competitive performance against a baseline using explicit AlphaFold3-derived coordinates (recall 0.726 vs. 0.734, AUC 0.843 vs. 0.898) with an 11.6× reduction in features. The most predictive feature (Triangle Count) directly reflects local residue packing density, a physical property typically requiring 3D coordinates. That the same encoding principles capture structurally meaningful information across both proteins and RNA strongly supports the interpretation of BioGraphX-RNA as a valid structure-informed proxy.

### 4.2 Cross-Species Transfer Is Limited and Sequence-Driven

The second question was whether structure-informed proxy representations capture evolutionarily conserved signals that aid cross-species generalization. Our blind cross-species test, which trained exclusively on human data and evaluated on mouse data, provides a clear answer: the biophysical graph features do not improve cross-species transfer. The RiNALMo-only baseline and the graph-only model yield near-identical macro AUROC values, both close to chance. The only compartment with consistent above-chance signal is the nucleus. These findings underscore that cross-species RNA localization prediction remains an open challenge, and that future progress will likely require species-specific training or the integration of phylogenetic information into the encoding.

### 4.3 lncRNA performance reflects inherent task difficulty

The relatively low macro-AUROC for lncRNA (0.5600 ± 0.0191 for the gated fusion model) raises the question of whether this reflects insufficient model capacity or a fundamental ceiling for sequence- and structure-based prediction. The ablation study (Supplementary Table S8) indicates that no configuration substantially outperforms the others: the RiNALMo-only baseline achieves 0.5889 ± 0.0247, simple concatenation 0.5654 ± 0.0049, and the graph-only model 0.5408 ± 0.0074. That the language model alone performs at least as well as any hybrid model suggests that the available sequence and biophysical features are approaching the information limit for this task, rather than the model being under-parameterized. This is consistent with the known biology of lncRNAs: their localization is governed by highly combinatorial and context-dependent mechanisms, including cell-type-specific expression, alternative splicing, and interactions with RNA-binding proteins that are not fully captured by primary sequence or local structure alone.

Future improvements in lncRNA localization prediction will likely require integrating additional data modalities, such as tissue-specific expression profiles or experimentally determined protein-binding sites.

### 4.4 BioGraphX-RNA Generates Testable Hypotheses

The third and arguably most important question was whether there existed a possibility of deriving interpretable predictions that would provide mechanistic insights into RNA trafficking based on explicit biochemical constraints. Our multi-faceted explainability toolkit generates testable hypotheses about the structural basis of subcellular localization, providing a starting point for future experimental validation.

### 4.5 Gating Reveals RNA-Type-Specific Structural Dependencies

The gating mechanism provides a systems-level view of modality reliance (Fig. 3). For mRNA, the model consistently assigns a higher weight to sequence embeddings, consistent with the view that mRNA localization is primarily driven by linear motifs recognized by RNA-binding proteins. For miRNA, the gate values approach an equal balance between sequence and structure, reflecting the absolute structural requirements of miRNA biology, from precursor hairpin recognition to RISC loading. For lncRNA, the gate distribution is intermediate and broader, capturing the functional diversity of this class. These patterns demonstrate that the gated architecture, while not improving accuracy, dynamically adjusts to each RNA type and offers interpretable hypotheses about the relative importance of sequence versus structural information.

### 4.6 SHAP Analysis Reveals Associative Patterns Across Compartments

The SHAP associations described in Section 3.5 are hypothesis-generating: they do not establish causal mechanisms but point to structural correlates that warrant experimental follow-up. Two patterns emerge consistently. First, GC periodicity (GC Autocorrelation Lag1) appears as a top positive feature for nuclear and ribosomal localization, suggesting that regularly spaced GC elements, rather than absolute GC content, are a recurring motif in localization signals. Second, structural accessibility (high Backbone Ratio, low local connectivity) correlates with exosome targeting across both mRNA and miRNA, while the absence of protective hybrid structures is a common repeller. These patterns align with the known biology of nuclear retention signals and exosome-mediated decay, but their confirmation requires independent experiments such as RBP-binding or CLIP-seq data.

### 4.7 Implications for RNA Biology and Precision Medicine

The mechanistic understanding gained through BioGraphX-RNA extends beyond the predictive power of the approach. By pinpointing particular structural elements that correlate with each compartment, our model produces testable hypotheses. For instance, the prediction that exosome targeting is mediated by structural accessibility rather than AREs could be confirmed by mutagenesis experiments that abrogate protective structures while maintaining sequence motifs. Likewise, the role of GC periodicity in nuclear retention implies that engineered RNAs could be designed with particular spacing patterns to regulate their cellular localization.

Within the realm of precision medicine, where RNA localization disorders are increasingly recognized as a cause of human disease, from cancer to neurodegenerative diseases , BioGraphX-RNA represents a resource for uncovering structurally encoded signals that could be compromised by disease-causing mutations.

### 4.8 Universality of the BioGraphX Encoding Paradigm

The BioGraphX framework is not limited to any particular domain. It is a universal encoding strategy applicable to any linear biological polymer. The core algorithm remains the same for different molecular families; only the rules differ to reflect the unique physicochemical context of each molecule type. We previously demonstrated this concept for amino acid sequences in proteins and proposed its extension to nucleic acids. In this study, we verify this hypothesis for RNA by simply replacing the protein interaction rules with Watson-Crick base pairing, stacking, and wobble rules. With BioGraphX now validated on both proteins and RNA across three distinct biological prediction tasks, the framework provides a unified, structure-informed encoding strategy whose extension to DNA or other domains is straightforward.

### 4.9 Limitations and Future Directions

Despite its strengths, BioGraphX-RNA also has some weaknesses that point to future improvements. Firstly, the graph-building approach uses linear sequence distance as a surrogate for physical proximity. Although this works well, it is still an approximation, and real three-dimensional structure, when known, might offer better interaction graphs. The current rapid progress in RNA structure prediction may soon allow the incorporation of predicted 3D coordinates into our approach, which could potentially offer better results.

Secondly, the current interaction rules are fixed and based on traditional biophysical knowledge. However, real RNA folding is a dynamic process, and RNA molecules have structures that exist as a mixture of different conformations and adapt to the cellular environment. Future updates might include in vivo RNA structural probing information to inform graph building or model conformational ensembles. Furthermore, the interaction types could be expanded to include RNA-protein interactions, which could potentially offer a more comprehensive view of localization mechanisms.

Third, although improved, the prediction of lncRNA is still a difficult task. The extreme length and functional diversity of lncRNAs might need more complex processing methods, such as using attention-based aggregation on window-level features or incorporating tissue-specific expression information. The distribution of gates for lncRNAs indicates that the heterogeneity of the classes is large; pre-classifying lncRNAs into functional or structural subtypes could potentially help.

Fourth, We did not perform a dedicated cross-RNA-family benchmark due to the lack of comprehensive family annotations in current datasets. This remains a valuable direction for future work.

Fifth, while we used the DeepLocRNA benchmark for comparison with prior work, direct evaluation against recent methods such as RNALoc-LM and LGLoc was not feasible due to differences in dataset partitions and the computational cost of retraining. Future benchmarking on standardized resources such as RNALocate v3.0 would enable more comprehensive comparisons.

Sixth, the SHAP-derived associations reported are model-specific and correlational; validation against external signals such as RBP-binding or CLIP data remains an important direction for future experimental follow-up.

## 5 Conclusion

In this work, we propose BioGraphX-RNA, a physicochemical graph representation framework that uses biophysical rules to encode structural priors for explainable subcellular localization prediction.

By representing nucleotide sequences as multiscale interaction graphs based on biophysical models, and incorporating frozen RiNALMo embeddings via an interpretable gated fusion, our model achieves competitive performance with DeepLocRNA, with substantial gains for miRNA. The gated fusion uniquely reveals the balance between sequence and structural signals for each RNA type.

Direct comparison with RNAfold-derived graphs, including a partition-function baseline that uses the full thermodynamic ensemble, confirms that BioGraphX-RNA captures structural signals beyond those available from folding-based representations, particularly for miRNA.

Cross-species evaluation on blind mouse data shows that transfer performance is limited. Only nuclear retention signals exhibit modest, above-chance conservation.

Most notably, BioGraphX-RNA is an interpretable model. Gating analysis reveals RNA-type-specific modality reliance, and SHAP analysis points to candidate biophysical correlates that warrant experimental follow-up, though these are hypothesis-generating. These results establish a systems-level trade-off between structural stability and flexibility in RNA trafficking.

With only 2.05M trainable parameters and a total end-to-end compute of approximately 10 GPU-hours, the model adheres to Green AI principles while incorporating multi-scale structural determinants of RNA fate decisions.

In summary, BioGraphX-RNA shows that incorporating biophysical constraints into graph-based encodings enables accurate and interpretable predictions for structured RNAs, advancing structure-aware RNA biology and providing a foundation for precision medicine.

## Declarations

### Consent for publication

Not applicable.

### Ethics Approval and Consent to Participate

Not applicable.

## Acknowledgments

The author acknowledges the use of publicly accessible computational resources provided through the Kaggle platform for conducting the experiments in this study.

## Author contributions statement

A.S. conceived the study, developed the methodology, implemented the algorithms, conducted experiments, analyzed the data, and wrote the manuscript.

## Data Availability

The DeepLocRNA dataset analyzed in this study is publicly available from the original publication by Wang et al., “DeepLocRNA: an interpretable deep learning model for predicting RNA subcellular localization with domain-specific transfer-learning”, Bioinformatics, 2024, 40(2): btae065. https://doi.org/10.1093/bioinformatics/btae065.

The source code for BioGraphX-RNA is openly available on GitHub at: (https://github.com/AbubakarSaeed/BioGraphX-RNA).

## References

Abaturov, A. E. and Babych, V. L. (2021). Localization and translocation of mature mirnas. Child’s Health, 16(7):498–507.

Akbari Rokn Abadi, S., Shahbakhsh, A., and Koohi, S. (2025). Lgloc as a new language model-driven graph neural network for mrna localization. Scientific Reports, 15:18709.

Alejo, J. L., Girodat, D., Hammerling, M. J., Willi, J. A., Jewett, M. C., Engelhart, A. E., and Adamala, K. P. (2024). Alternate conformational trajectories in ribosome translocation. PLoS Computational Biology, 20(8):e1012319.

Baleriola, J., Walker, C. A., Jean, Y. Y., et al. (2014). Axonally synthesized atf4 transmits a neurodegenerative signal across brain regions. Cell, 158:1159–1172.

Bassell, G. J. and Warren, S. T. (2008). Fragile x syndrome: loss of local mrna regulation alters synaptic development and function. Neuron, 60:201–214.

Batista, L. F. Z., Pech, M. F., Zhong, F. L., et al. (2011). Telomere shortening and loss of self-renewal in dyskeratosis congenita induced pluripotent stem cells. Nature, 474:399–402.

Bauman, J. G. J., Wiegant, J., Borst, P., and van Duijn, P. (1980). A new method for fluorescence microscopical localization of specific dna sequences by in situ hybridization of fluorochrome-labelled rna. Experimental Cell Research, 128(2):485–490.

Bernstein, E., Caudy, A. A., Hammond, S. M., and Hannon, G. J. (2001). Role for a bidentate ribonuclease in the initiation step of rna interference. Nature, 409(6818):363–366.

Bugai, A., Hohmann, U., Lorenzo, A., Graf, M., Fin, L., Rouviére, J. O., Tirian, L., Dou, Y., Polák, P., Johnsen, D., Jakobsen, L., Andersen, J. S., Brennecke, J., Plaschka, C., and Jensen, T. H. (2025). Molecular basis of polyadenylated rna fate determination in the nucleus. bioRxiv.

Carlevaro-Fita, J. and Johnson, R. (2019). Global positioning system: Understanding long noncoding rnas through subcellular localization. Molecular Cell, 73(5):869–883.

Chen, C.-Y., Gherzi, R., Ong, S.-E., Chan, E. L., Raijmakers, R., Pruijn, G. J., Stoecklin, G., Moroni, C., Mann, M., and Karin, M. (2001). Au binding proteins recruit the exosome to degrade are-containing mrnas. Cell, 107(4):451–464.

Chen, S. and Collart, M. A. (2024). Membrane-associated mrnas: A post-transcriptional pathway for fine-tuning gene expression. Journal of Molecular Biology, 436:168579.

Crick, F. H. C. (1966). Codon–anticodon pairing: The wobble hypothesis. Journal of Molecular Biology, 19(2):548–555.

Dictenberg, J. B., Swanger, S. A., Antar, L. N., et al. (2008). A direct role for fmrp in activity-dependent dendritic mrna transport links filopodial-spine morphogenesis to fragile x syndrome. Developmental Cell, 14:926–939.

Didiot, M.-C., Ferguson, C. M., Ly, S., et al. (2018). Nuclear localization of huntingtin mrna is specific to cells of neuronal origin. Cell Reports, 24:2553–2560.e5.

Doudna, J. A. and Cech, T. R. (2002). The chemical repertoire of natural ribozymes. Nature, 418(6894):222–228.

Fu, L., Niu, B., Zhu, Z., Wu, S., and Li, W. (2012). Cd-hit: accelerated for clustering the next-generation sequencing data. Bioinformatics, 28(23):3150–3152.

Garcia-Martin, R. et al. (2022). Microrna sequence codes for small extracellular vesicle release and cellular retention. Nature, 601(7893):446–451.

Grishok, A., Pasquinelli, A. E., Conte, D., Li, N., Parrish, S., Ha, I. K., Baillie, D. L., Fire, A., Ruvkun, G., and Mello, C. C. (2001). Genes and mechanisms related to rna interference regulate expression of the small temporal rnas that control c. elegans developmental timing. Cell, 106(1):23–34.

Guo, C. J. et al. (2020). Distinct processing of lncrnas contributes to non-conserved functions in stem cells. Cell, 181(3):621–636.e22.

Guo, Y. et al. (2015). Rnaseq by total rna library identifies additional rnas compared to poly(a) rna library. BioMed Research International, 2015:862130.

Helena-Bueno, K., Brown, A., Smith, J., et al. (2025). Structurally heterogeneous ribosomes cooperate in protein synthesis in bacterial cells. Nature Communications, 16(1):2751.

Huang, Z. et al. (2025). Exosome mirna sorting controlled by rna-binding protein-motif interactions. Extracellular Vesicles and Circulating Nucleic Acids, 6(3):475–503.

Ivy, A. S., Rex, C. S., Chen, Y., et al. (2010). Hippocampal dysfunction and cognitive impairments provoked by chronic early-life stress involve excessive activation of crh receptors. Journal of Neuroscience, 30:13005–13015.

Jan, C. H. et al. (2014). Single-molecule quantification of translation-dependent association of mrnas with the endoplasmic reticulum. Cell Reports.

Jao, L.-E., Akef, A., Wente, S. R., et al. (2017). A role for gle1, a regulator of dead-box rna helicases, at centrosomes and basal bodies. Molecular Biology of the Cell, 28:120–127.

Keidel, A., Kögel, A., Reichelt, P., Kowalinski, E., Schäfer, I. B., and Conti, E. (2023). Concerted structural rearrangements enable rna channeling into the cytoplasmic ski238-ski7-exosome assembly. Molecular Cell, 83(22):4093–4105.e7.

Kim, S. et al. (2025). Exogenous rna surveillance by proton-sensing trim25. Science.

Leontis, N. B. and Westhof, E. (2001). Geometric nomenclature and classification of rna base pairs. RNA, 7(4):499–512.

Leucci, E., Vendramin, R., Spinazzi, M., et al. (2016). Melanoma addiction to the long non-coding rna sammson. Nature, 531:518–522.

Lin, W.-J., Duffy, A., and Chen, C.-Y. (2007). Localization of au-rich element-containing mrna in cytoplasmic granules containing exosome subunits. Journal of Biological Chemistry, 282(27):19958–19968.

Lorenz, R., Bernhart, S. H., Höner zu Siederdissen, C., Tafer, H., Flamm, C., Stadler, P. F., and Hofacker, I. L. (2011). Viennarna package 2.0. Algorithms for Molecular Biology, 6(1):26.

Lubas, M. et al. (2015). The human nuclear exosome targeting complex is loaded onto newly synthesized rna to direct early ribonucleolysis. Cell Reports, 10(2):178–192.

Lubelsky, Y. and Ulitsky, I. (2018). Sequences enriched in alu repeats drive nuclear localization of long rnas in human cells. Nature, 555(7694):107–111.

Ma, W. and Mayr, C. (2018). A membraneless organelle associated with the endoplasmic reticulum enables 3 utrmediated protein-protein interactions. Cell, 175(6):1492–1506.e19.

Ma, W., Zhen, G., Xie, W., and Mayr, C. (2021). In vivo reconstitution finds multivalent rna–rna interactions as drivers of mesh-like condensates. eLife, 10:e64252.

Mathews, D. H., Disney, M. D., Childs, J. L., Schroeder, S. J., Zuker, M., and Turner, D. H. (2004). Incorporating chemical modification constraints into a dynamic programming algorithm for prediction of rna secondary structure. Proceedings of the National Academy of Sciences, 101(19):7287–7292.

Neelamraju, Y., Gonzalez-Perez, A., Bhat-Nakshatri, P., et al. (2018). Mutational landscape of rna-binding proteins in human cancers. RNA Biology, 15:115–129.

Noh, J. H., Kim, K. M., McClusky, W. G., Abdelmohsen, K., and Gorospe, M. (2018). Cytoplasmic functions of long noncoding rnas. Wiley Interdisciplinary Reviews: RNA, 9(3):e1471.

Nousiainen, H. O., Kestilä, M., Pakkasjärvi, N., et al. (2008). Mutations in mrna export mediator gle1 result in a fetal motoneuron disease. Nature Genetics, 40:155–157.

Okamura, M., Yamanaka, Y., Shigemoto, M., et al. (2019). Depletion of mrna export regulator dbp5/ddx19, gle1 or ippk that is a key enzyme for the production of ip6, resulting in differentially altered cytoplasmic mrna expression and specific cell defect. PLoS One, 14:e0197165.

Otto, G. (2019). Protein trafficking through tiger domains. Nature Reviews Molecular Cell Biology, 20:3.

Panda, S., Setia, M., Kaur, N., et al. (2018). Noncoding rna ginir functions as an oncogene by associating with centrosomal proteins. PLoS Biology, 16:e2004204.

Penić, R. J., Vla šić, T., Huber, R. G., Wan, Y., and Šikić, M. (2025). Rinalmo: general-purpose rna language models can generalize well on structure prediction tasks. Nature Communications, 16:5671.

Saeed, A. and Abbas, W. (2026). Biographx: bridging the sequence-structure gap via physicochemical graph encoding for interpretable subcellular localization prediction. Bioinformatics Advances, 6(1):vbag181.

Saenger, W. (1984). Principles of Nucleic Acid Structure. Springer-Verlag, New York.

Sehgal, A., Briggs, J., Rinehart-Kim, J., Basso, J., and Bos, T. J. (2000). The chicken c-jun 5′ untranslated region directs translation by internal initiation. Oncogene, 19(24):2836–2845.

Sun, X. L. and Antony, A. C. (1996). Evidence that a specific interaction between an 18-base cis-element in the 5′-untranslated region of human folate receptor-α mrna and a 46-kda cytosolic trans-factor is critical for translation. Journal of Biological Chemistry.

Tang, W. and Tseng, H. (1999). A gc-rich sequence within the 5′ untranslated region of human basonuclin mrna inhibits its translation. Gene, 237(1):35–44.

Tinoco, I. J., Borer, P. N., Dengler, B., Levine, M. D., Uhlenbeck, O. C., Crothers, D. M., and Gralla, J. (1973). Improved estimation of secondary structure in ribonucleic acids. Nature New Biology, 246:40–41.

Vigh, M. L., Bressendorff, S., Thieffry, A., Arribas-Hernández, L., and Brodersen, P. (2022). Nuclear and cytoplasmic rna exosomes and pelota1 prevent mirna-induced secondary sirna production in arabidopsis. Nucleic Acids Research, 50(3):1396–1415.

Vihinen, M. (2021). Systematics for types and effects of rna variations. RNA Biology, 18(4):481–498.

Villarroya-Beltri, C. et al. (2013). Sumoylated hnrnpa2b1 controls the sorting of mirnas into exosomes through binding to specific motifs. Nature Communications, 4:2980.

Wang, J., Horlacher, M., Cheng, L., and Winther, O. (2024a). Deeplocrna: an interpretable deep learning model for predicting rna subcellular localization with domain-specific transfer-learning. Bioinformatics, 40(2):btae065.

Wang, N., Bian, J., Li, Y., Li, X., Xiong, H., et al. (2024b). Multi-purpose rna language modelling with motif-aware pretraining and type-guided fine-tuning. Nature Machine Intelligence, 6:548–557.

Watson, J. D. and Crick, F. H. C. (1953). Molecular structure of nucleic acids: A structure for deoxyribose nucleic acid. Nature, 171(4356):737–738.

Wu, L., Wang, L., Hu, S., Tang, G., Chen, J., Yi, Y., Xie, H., Lin, J., Wang, M., Wang, D., Yang, B., and Huang, Y. (2025). Rnalocate v3.0: Advancing the repository of rna subcellular localization with dynamic analysis and prediction. Nucleic Acids Research, 53(D1):D284–D292.

Yao, J., Tu, Y., Shen, C., Zhou, Q., Xiao, H., Jia, D., and Sun, Q. (2021). Nuclear import receptors and hnrnpk mediates nuclear import and stress granule localization of sirloin. Cellular and Molecular Life Sciences, 78(23):7617–7633.

Zeng, M., Zhang, X., Li, Y., Lu, C., Yin, R., Guo, F., and Li, M. (2025). Rnaloc-lm: Rna subcellular localization prediction using pre-trained rna language model. Bioinformatics, 41(4):btaf127.

Zhang, B., Babu, K. R., Lim, C. Y., et al. (2020). A comprehensive expression landscape of rna-binding proteins (rbps) across 16 human cancer types. RNA Biology, 17:211–226.

Zhang, H., Kolb, F. A., Brondani, V., Billy, E., and Filipowicz, W. (2004). Single processing center models for human dicer and bacterial rnase iii. Nature, 431(7010):248–252.

Zuckerman, B. and Ulitsky, I. (2019). Predictive models of subcellular localization of long rnas. RNA, 25(5):557–572.

