## Supplementary Material for "BioGraphX-RNA: A Universal Physicochemical Graph Encoding for Interpretable RNA Subcellular Localization Prediction"

### Supplementary Material S1: Complete BioGraphX-RNA Feature Descriptions (149 Features)

This supplementary document provides a comprehensive description of all 149 features extracted by the BioGraphX-RNA framework. The features are derived from an RNA-specific biochemically-calibrated constraint graph constructed directly from nucleotide sequences. The framework's multi-scale approach captures RNA topology, base pairing networks, physicochemical properties, and structural frustration to create a structure-informed proxy without requiring 3D coordinates.

#### 0.1 1. Topological RNA Graph Features (65 features)

*Description: Network architecture metrics derived from the RNA biochemical interaction graph, applying concepts from graph theory to model nucleotide relationships, including canonical Watson-Crick pairing, wobble pairs, base stacking, and backbone connectivity.*

Table 1: Topological RNA Graph Features (65 total)

| **Feature Name** | **Description** | **Biological Interpretation / Basis** |
|---|---|---|
| Node_Count | Number of nucleotides (vertices) in the graph. | RNA length. |
| Edge_Count | Number of RNA interactions (edges) in the graph. | Overall structural connectivity. |
| Graph_Density | Ratio of actual edges to possible edges. | Structural compactness. |
| Mean_Degree | Average number of interactions per nucleotide. | Average nucleotide connectivity. |
| Degree_Std | Standard deviation of degree distribution. | Heterogeneity in local structure. |
| Max_Degree | Maximum number of interactions for any nucleotide. | Identification of structural hubs (junctions). |
| Degree_Q1 | First quartile of degree distribution. | Lower bound of connectivity. |
| Degree_Median | Median of degree distribution. | Central tendency of connectivity. |
| Degree_Q3 | Third quartile of degree distribution. | Upper bound of connectivity. |
| Mean_Weighted_Degree | Average edge-weighted degree (by interaction strength). | Strength of interactions per nucleotide. |
| Weighted_Degree_Std | Variability in weighted degree. | Diversity in interaction strengths. |
| Max_Weighted_Degree | Maximum weighted degree. | Nucleotide with strongest total interactions. |
| Canonical_WC_Ratio | Proportion of Watson-Crick base pairing edges. | Core secondary structure stability Turner and Mathews (2009). |

Table 1: Topological RNA Graph Features (65 total) - continued

| Feature Name | Description | Biological Interpretation / Basis |
|---|---|---|
| Wobble_Pair_Ratio | Proportion of G-U wobble base pairing edges. | Flexible base pairing, tRNA structure. |
| Non_Canonical_Ratio | Proportion of non-canonical base pairing edges. | Tertiary structure interactions, ribozymes. |
| Base_Stacking_Ratio | Proportion of base stacking interaction edges. | Helical stabilization, thermodynamic stability Yakovchuk et al. (2006). |
| Backbone_Ratio | Proportion of phosphate backbone edges. | Chain connectivity and flexibility. |
| Mean_Interaction_Energy | Average interaction energy estimate (kcal/mol). | Overall structural stability. |
| Energy_Std | Standard deviation of interaction energies. | Heterogeneity in interaction strengths. |
| Min_Energy | Most negative (strongest) interaction energy. | Strongest stabilizing interactions. |
| Total_Stabilization | Sum of all interaction energies. | Total estimated folding stability. |
| Strong_Interactions_Ratio | Proportion of interactions with energy $<$ -1.0 kcal/mol. | Fraction of strong stabilizing forces. |
| Mean_Betweenness | Average betweenness centrality. | Importance in structural communication Brandes (2001). |
| Betweenness_Std | Standard deviation of betweenness. | Heterogeneity in pathway importance. |
| Max_Betweenness | Maximum betweenness centrality. | Critical junction nucleotides. |
| Mean_Closeness | Average closeness centrality (cutoff=10). | Average proximity in local structure. |
| Closeness_Std | Standard deviation of closeness (cutoff=10). | Heterogeneity in proximity. |
| Max_Closeness | Maximum closeness centrality (cutoff=10). | Centrally located nucleotides. |
| Min_Closeness | Minimum closeness centrality (cutoff=10). | Remotely located nucleotides. |
| Mean_Eigenvector | Average eigenvector centrality. | Influence in structural neighborhoods. |
| Eigenvector_Std | Standard deviation of eigenvector centrality. | Heterogeneity in neighborhood influence. |
| Community_Count | Number of detected communities (structural domains). | Structural domain organization. |
| Modularity | Quality of community partition. | Strength of domain separation. |
| Mean_Community_Size | Average community size. | Typical structural domain size. |
| Community_Size_Std | Standard deviation of community sizes. | Heterogeneity in domain sizes. |
| Max_Community_Size | Largest community size. | Major structural domain. |
| IntraCommunity_Edge_Ratio | Ratio of edges within communities. | Internal domain connectivity. |
| Five_Prime_Connectivity | Edge density in 5' region (first 30 nucleotides). | 5' UTR structural compactness. |
| Five_Prime_GC | GC content in 5' region. | 5' structural stability (translation initiation). |
| Five_Prime_Strong_Edges | Ratio of strong edges in 5' region. | Stable structural elements near start. |
| Three_Prime_Connectivity | Edge density in 3' region (last 30 nucleotides). | 3' UTR structural compactness. |
| Three_Prime_AU | AU content in 3' region. | 3' regulatory element flexibility. |
| Three_Prime_Terminal_Degree | Degree of the last nucleotide. | 3' end accessibility (polyA tail). |
| GC_Cluster_Connectivity | Edge density in GC-rich positions. | Connectivity of stable structural regions. |

Table 1: Topological RNA Graph Features (65 total) - continued

| Feature Name | Description | Biological Interpretation / Basis |
|---|---|---|
| GC_Cluster_Count | Number of GC-rich clusters. | Multiple stable structural elements. |
| Max_GC_Cluster_Size | Size of largest GC-rich cluster. | Major stable structural domain. |
| AU_Cluster_Connectivity | Edge density in AU-rich positions. | Connectivity of flexible/unstructured regions. |
| AU_Cluster_Count | Number of AU-rich clusters. | Multiple flexible regulatory elements. |
| Max_AU_Cluster_Size | Size of largest AU-rich cluster. | Major flexible region. |
| Avg_Shortest_Path | Average shortest path length (LCC). | Global structural communication efficiency. |
| Graph_Diameter | Longest shortest path (LCC). | Maximum structural distance. |
| Global_Efficiency | Inverse of average shortest path length. | Structural communication efficiency. |
| Local_Efficiency | Average efficiency of local neighborhoods. | Local structural robustness Brandes (2001). |
| Path_Length_Std | Standard deviation of sampled paths. | Heterogeneity in structural distances. |
| Max_Path_Length | Maximum sampled shortest path. | Extreme structural distance. |
| Clustering_Coefficient | Average local clustering coefficient. | Local structural cohesiveness. |
| Assortativity | Degree correlation coefficient. | Homophily in connectivity. |
| Strong_Interaction_Ratio | Proportion of interactions with weight >0.7. | Fraction of strong interactions. |
| Mean_Energy | Average interaction energy. | Typical interaction strength. |
| Min_Energy_Overall | Minimum (strongest) interaction energy. | Strongest stabilizing interaction. |
| Stabilization_Per_Nucleotide | Total stabilization normalized by length. | Average structural contribution per nt. |
| Hybrid_Edge_Ratio | Proportion of hybrid interaction edges. | Interaction complexity. |
| Mean_Edge_Weight | Average edge weight. | Average interaction strength. |
| Edge_Weight_Std | Standard deviation of edge weights. | Heterogeneity in strengths. |
| Max_Edge_Weight | Maximum edge weight. | Strongest individual interaction. |

#### 0.2 2. RNA Hybrid Interaction Features (15 features)

*Description: Co-occurrence patterns of RNA interactions where multiple interaction types occur simultaneously between the same nucleotide pair. These synergistic combinations represent specialized structural motifs critical for function.*

Table 2: RNA Hybrid Interaction Features (15 total)

| Feature Name | Description | Biological Interpretation |
|---|---|---|
| WC_Stacking_Hybrid | Frequency of Watson-Crick pairing + base stacking. | Stable helical regions, canonical stems. |
| Wobble_Stacking_Hybrid | Frequency of wobble pairing + base stacking. | Flexible helices, tRNA acceptor stems. |
| Stacking_Backbone_Hybrid | Frequency of base stacking + backbone connectivity. | Tight helical turns, loop regions. |
| Five_Prime_Hybrid_Density | Hybrid density in 5' region (first 30 nt). | Complex 5' regulatory structures. |
| Three_Prime_Hybrid_Density | Hybrid density in 3' region (last 30 nt). | Complex 3' regulatory structures. |
| GC_Region_Hybrid_Density | Hybrid density in GC-rich positions. | Specialized stable structural motifs. |
| AU_Region_Hybrid_Density | Hybrid density in AU-rich positions. | Specialized flexible regulatory motifs. |
| Hybrid_Edge_Ratio_Detailed | Proportion of edges that are hybrid. | Overall interaction complexity. |

Table 2: RNA Hybrid Interaction Features (15 total) - continued

| Feature Name | Description | Biological Interpretation |
|---|---|---|
| Hybrid_Subgraph_Density | Density of subgraph of hybrid edges. | Compactness of hybrid clusters. |
| Hybrid_Cluster_Count | Connected components in hybrid subgraph. | Modularity of hybrid interactions. |
| Mean_Hybrid_Energy | Average energy of hybrid interactions. | Stability of synergistic motifs. |
| Min_Hybrid_Energy | Minimum (strongest) hybrid energy. | Strongest synergistic interaction. |
| Strong_Hybrid_Ratio | Proportion of hybrid edges with energy $<$ -1.0. | Fraction of strong synergistic motifs. |
| Hybrid_Diversity | Shannon entropy of hybrid types. | Diversity of hybrid interactions. |
| Hybrid_Connectivity | Average degree in hybrid subgraph. | Connectivity of hybrid network. |

### 0.3 3. RNA Knowledge-Guided Profile Features (27 features)

*Description: Sequence-based localization signals derived from known RNA motifs and hybrid scores, informed by databases of experimentally validated RNA localization signals and subcellular distribution patterns.*

Table 3: RNA Knowledge-Guided Profile Features (27 total)

| Feature Name | Description | Biological Interpretation |
|---|---|---|
| Profile_Nucleus_Motif | Nuclear localization motif score. | G/A-rich clusters, RRKY motifs, nuclear retention signals Zhang et al. (2020). |
| Profile_Nucleus_Hybrid | Hybrid-based nuclear compatibility. | Structural compatibility with nuclear environment. |
| Profile_Nucleus_GC_Compat | GC content compatibility with nucleus. | Nuclear RNAs typically 40-60% GC. |
| Profile_Exosome_Motif | Exosome targeting/degradation score. | AU-rich elements (AREs), UUAUUUAUU motifs Chen and Shyu (1995). |
| Profile_Exosome_Hybrid | Hybrid-based exosome compatibility. | Structural accessibility to exosome complex. |
| Profile_Exosome_GC_Compat | GC content compatibility with exosome. | Degradation-prone RNAs often AU-rich (<50% GC). |
| Profile_Cytosol_Motif | Cytosolic localization score. | Balanced nucleotide composition, no strong signals. |
| Profile_Cytosol_Hybrid | Hybrid-based cytosolic compatibility. | Compatibility with cytosolic environment. |
| Profile_Cytosol_GC_Compat | GC content compatibility with cytosol. | Balanced GC content (40-60%). |
| Profile_Cytoplasm_Motif | Cytoplasmic localization score. | A/U-rich clusters, mRNA-like composition. |
| Profile_Cytoplasm_Hybrid | Hybrid-based cytoplasmic compatibility. | Compatibility with cytoplasmic environment. |
| Profile_Cytoplasm_GC_Compat | GC content compatibility with cytoplasm. | Moderate GC content (35-55%). |
| Profile_Ribosome_Motif | Ribosome association/translation score. | Shine-Dalgarno-like, Kozak context, purine-rich. |
| Profile_Ribosome_Hybrid | Hybrid-based ribosome compatibility. | Structural compatibility with ribosome binding. |
| Profile_Ribosome_GC_Compat | GC content compatibility with ribosome. | Ribosomal RNAs GC-rich (>50%). |
| Profile_Membrane_Motif | Membrane association score. | U-rich clusters, hydrophobic k-mers Yan et al. (2016). |

Table 3: RNA Knowledge-Guided Profile Features (27 total) - continued

| **Feature Name** | **Description** | **Biological Interpretation** |
|---|---|---|
| Profile_Membrane_Hybrid | Hybrid-based membrane compatibility. | Compatibility with membrane environment. |
| Profile_Membrane_GC_Compat | GC content compatibility with membrane. | Membrane-associated RNAs often AU-rich. |
| Profile_Endoplasmic_reticulum_Motif | ER targeting score. | SRP binding signals, ER retention motifs. |
| Profile_Endoplasmic_reticulum_Hybrid | Hybrid-based ER compatibility. | Compatibility with ER lumen/membrane. |
| Profile_Endoplasmic_reticulum_GC_Compat | GC content compatibility with ER. | Moderate GC content (30-60%). |
| Profile_Microvesicles_Motif | Microvesicle/exosome packaging score. | U-rich clusters, specific export motifs Valadi et al. (2007). |
| Profile_Microvesicles_Hybrid | Hybrid-based microvesicle compatibility. | Compatibility with vesicle packaging. |
| Profile_Microvesicles_GC_Compat | GC content compatibility with microvesicles. | Secreted RNAs often AU-rich (<50% GC). |
| Profile_Mitochondrion_Motif | Mitochondrial targeting score. | A-rich 5' regions, mito-targeting motifs. |
| Profile_Mitochondrion_Hybrid | Hybrid-based mitochondrial compatibility. | Compatibility with mitochondrial import. |
| Profile_Mitochondrion_GC_Compat | GC content compatibility with mitochondrion. | Mitochondrial RNAs often GC-rich (>50%). |

#### 0.4 4. Global RNA Physicochemical Features (25 features)

*Description: Whole-molecule biophysical properties derived from nucleotide composition and sequence Freier et al. (1986); Mathews et al. (1999), reflecting thermodynamic constraints important for RNA folding and stability.*

Table 4: Global RNA Physicochemical Features (25 total)

| **Feature Name** | **Description** | **Biological Interpretation** |
|---|---|---|
| GC_Content | Fraction of G and C nucleotides. | Primary determinant of thermal stability. |
| AU_Content | Fraction of A and U nucleotides. | Correlated with flexibility and instability. |
| A_Frequency | Fraction of adenine. | Base composition. |
| U_Frequency | Fraction of uracil. | Base composition. |
| G_Frequency | Fraction of guanine. | Base composition. |
| C_Frequency | Fraction of cytosine. | Base composition. |
| GC_Skew | (G-C)/(G+C). | Strand asymmetry, transcription effects. |
| AU_Skew | (A-U)/(A+U). | Compositional strand bias. |
| Shannon_Entropy | Sequence complexity (nucleotide distribution). | Information content, sequence variability Schneider and Stephens (1990). |

Table 4: Global RNA Physicochemical Features (25 total) - continued

| Feature Name | Description | Biological Interpretation |
|---|---|---|
| MFE_Per_Nucleotide | Approximate minimum free energy per nucleotide. | Folding stability normalized by length Turner and Mathews (2009). |
| AA_Frequency | Dinucleotide frequency (AA). | Nearest-neighbor stacking propensity. |
| UU_Frequency | Dinucleotide frequency (UU). | Nearest-neighbor stacking propensity. |
| GG_Frequency | Dinucleotide frequency (GG). | Nearest-neighbor stacking propensity. |
| CC_Frequency | Dinucleotide frequency (CC). | Nearest-neighbor stacking propensity. |
| AU_Frequency | Dinucleotide frequency (AU). | Nearest-neighbor stacking propensity. |
| UA_Frequency | Dinucleotide frequency (UA). | Nearest-neighbor stacking propensity. |
| GC_Frequency | Dinucleotide frequency (GC). | Nearest-neighbor stacking propensity. |
| CG_Frequency | Dinucleotide frequency (CG). | Nearest-neighbor stacking propensity. |
| GU_Frequency | Dinucleotide frequency (GU). | Nearest-neighbor stacking propensity. |
| UG_Frequency | Dinucleotide frequency (UG). | Nearest-neighbor stacking propensity. |
| Mean_Pairing_Potential | Average base pairing potential (sliding window). | Local structural potential Hofacker et al. (1994). |
| Std_Pairing_Potential | Standard deviation of pairing potential. | Heterogeneity in structural potential. |
| Max_Pairing_Potential | Maximum pairing potential. | Regions with highest structural propensity. |
| Min_Pairing_Potential | Minimum pairing potential. | Regions with lowest structural propensity. |
| GC_Autocorrelation_Lag1 | GC content autocorrelation at lag 1. | Alternating GC patterns, structural periodicity. |

#### 0.5 5. RNA Frustration Features (17 features)

*Description: Per-nucleotide conflicting interaction energies derived from variance in constraint graph weights. These features capture biophysical conflicts that arise from competing base pairing possibilities, stacking vs. pairing trade-offs, and secondary structure constraints.*

Table 5: RNA Frustration Features (17 total)

| Feature Name | Description | Biological Interpretation |
|---|---|---|
| Frustration_5Prime_Mean | Mean frustration in 5' region (first 50 nucleotides). | 5' UTR structural conflicts vs. regulatory signals. |
| Frustration_3Prime_Mean | Mean frustration in 3' region (last 30 nucleotides). | 3' UTR structural conflicts vs. localization signals. |
| Frustration_Middle_Mean | Mean frustration in middle structural region. | Core folding constraint conflicts. |
| Frustration_GC_Mean | Mean frustration in GC-rich positions. | Conflicts in stable structural regions. |
| Frustration_GC_Variance | Variance of frustration in GC-rich positions. | Heterogeneity of conflicts in stable regions. |
| Frustration_AU_Mean | Mean frustration in AU-rich positions. | Conflicts in flexible/unstructured regions. |
| Frustration_AU_Variance | Variance of frustration in AU-rich positions. | Heterogeneity of conflicts in flexible regions. |
| Frustration_5vs3_Prime | Difference between 5' and 3' frustration. | Asymmetry in regulatory signal conflicts. |
| Frustration_StructuredVs Flexible | Difference between GC and AU frustration. | Conflict bias between structured and flexible regions. |

Table 5: RNA Frustration Features (17 total) - continued

| Feature Name | Description | Biological Interpretation |
|---|---|---|
| Frustration_Paired_Mean | Mean frustration in base-paired nucleotides. | Conflicts within paired regions. |
| Frustration_HotspotCount | Count of residues with frustration > mean + $1\sigma$. | Number of severe constraint conflict sites. |
| Frustration_HotspotDensity | Proportion of hotspots normalized by length. | Density of conflict sites. |
| Frustration_Motif_Mean | Mean frustration in predicted motif regions. | Conflicts overlapping regulatory signals. |
| Frustration_Motif_Max | Maximum frustration in motif regions. | Peak conflict in signal regions. |
| Frustration_NonMotif_Mean | Mean frustration in non-motif regions. | Background conflict level. |
| Frustration_LongRange_Mean | Mean frustration in long-range interactions (>20 nt apart). | Conflicts involving long-range tertiary contacts. |
| Frustration_Entropy | Shannon entropy of frustration distribution. | How localized vs. distributed conflicts are. |

### Supplementary Material S2: BioGraphX-RNA Feature Summary and SHAP Framework

Table 6 provides a summary of the five feature categories. The SHAP (SHapley Additive exPlanations) framework Lundberg and Lee (2017) was used to explain model predictions, providing theoretically sound quantification of each feature's contribution.

Table 6: Summary of BioGraphX-RNA Feature Categories (Total: 149 features)

| Category | # Features | Scale | Biological Insight |
|---|---|---|---|
| Topological RNA Graph | 65 | Nucleotide to molecule | RNA fold complexity, structural domain organization, network properties |
| RNA Hybrid Interaction | 15 | Pairwise interactions | Specialized structural motifs (helices, turns), synergistic base interactions |
| Knowledge-Guided Profile | 27 | Whole-molecule | Known RNA targeting signals + structural compatibility with compartments |
| Global Physicochemical | 25 | Whole-molecule | Thermodynamic constraints, folding stability, compositional biases |
| RNA Frustration | 17 | Per-nucleotide | Conflicting base pairing energies, structural compatibility filters |
| **Total** | **149** | **Multi-scale** | **Comprehensive sequence-structure-function representation** |

Table 7: Original vs Homology-Free Performance

| RNA Type | Metric | Original | Homology-free | $\Delta$ |
|---|---|---|---|---|
| mRNA | macro AUROC | $0.7575 \pm 0.0054$ | $0.7475 \pm 0.0045$ | $-0.0100$ |
| | macro AUPRC | $0.5870 \pm 0.0017$ | $0.5797 \pm 0.0024$ | $-0.0073$ |
| miRNA | macro AUROC | $0.9228 \pm 0.0137$ | $0.9193 \pm 0.0118$ | $-0.0035$ |
| | macro AUPRC | $0.7379 \pm 0.0125$ | $0.7299 \pm 0.0114$ | $-0.0080$ |
| lncRNA | macro AUROC | $0.5600 \pm 0.0191$ | $0.5648 \pm 0.0162$ | $+0.0048$ |
| | macro AUPRC | $0.3640 \pm 0.0104$ | $0.3622 \pm 0.0073$ | $-0.0018$ |

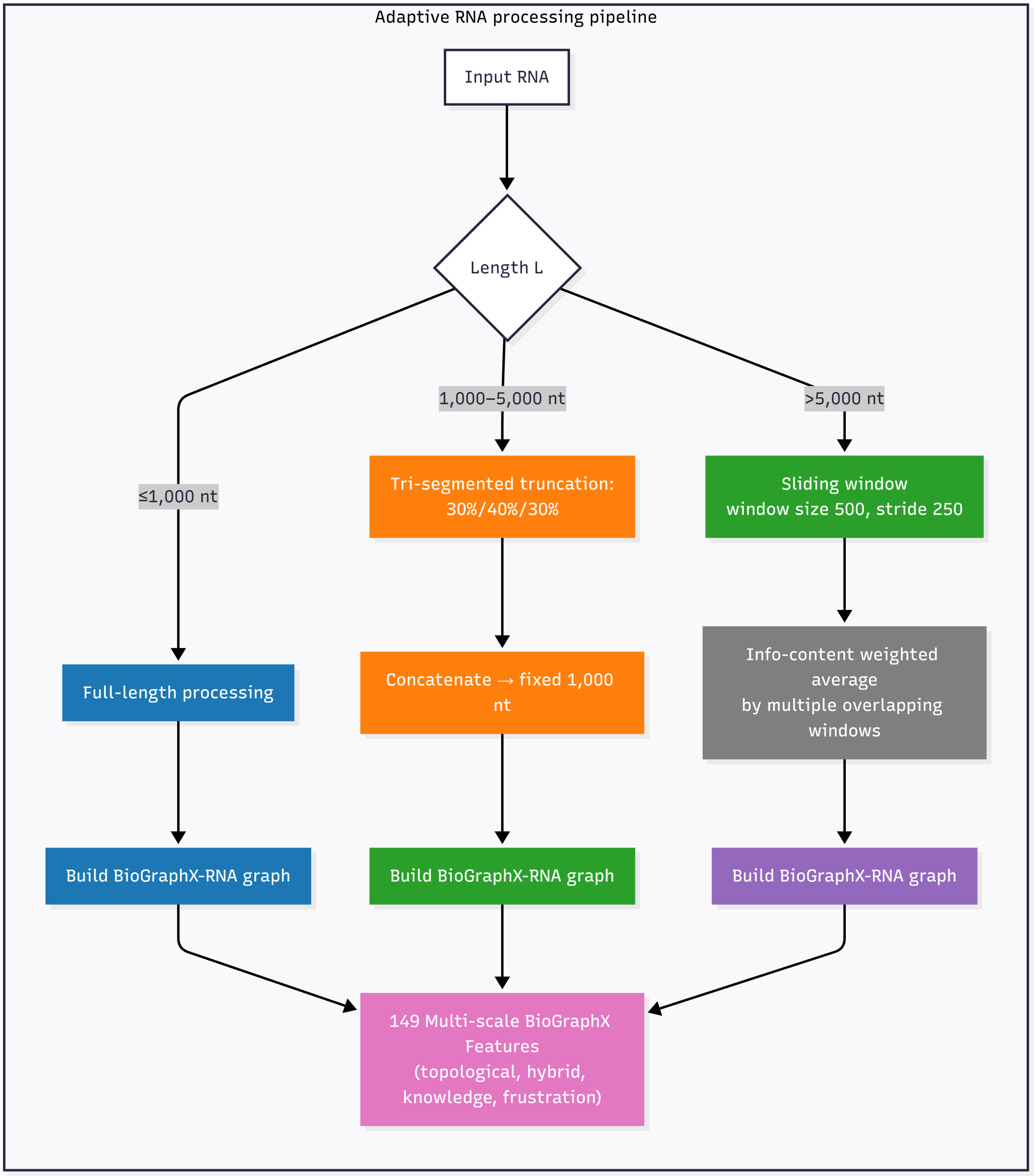


Figure 1: Adaptive RNA processing pipeline for short, medium, and long transcripts. Sequences are handled by full-length processing (short), tri-segmented truncation (medium), or sliding window with information-content-weighted averaging (long). Each pathway constructs a BioGraphX graph and extracts 149 multi-scale features for downstream classification.

Table 8: Ablation Study Results (Mean $\pm$ Standard Deviation)

| RNA Type | Model | Macro AUROC | Macro AUPRC |
|---|---|---|---|
| mRNA | RiNALMo-only | 0.7572 $\pm$ 0.0074 | **0.5910 $\pm$ 0.0026** |
| | Graph-only (MLP) | 0.6810 $\pm$ 0.0059 | 0.4924 $\pm$ 0.0011 |
| | Simple concatenation | **0.7581 $\pm$ 0.0028** | 0.5879 $\pm$ 0.0026 |
| | Gated fusion | 0.7575 $\pm$ 0.0054 | 0.5870 $\pm$ 0.0017 |
| miRNA | RiNALMo-only | 0.9157 $\pm$ 0.0069 | 0.7359 $\pm$ 0.0098 |
| | Graph-only (MLP) | **0.9396 $\pm$ 0.0045** | 0.7191 $\pm$ 0.0070 |
| | Simple concatenation | 0.9149 $\pm$ 0.0152 | 0.7293 $\pm$ 0.0109 |
| | Gated fusion | 0.9228 $\pm$ 0.0137 | **0.7379 $\pm$ 0.0125** |
| lncRNA | RiNALMo-only | **0.5889 $\pm$ 0.0247** | **0.3656 $\pm$ 0.0073** |
| | Graph-only (MLP) | 0.5408 $\pm$ 0.0074 | 0.3451 $\pm$ 0.0065 |
| | Simple concatenation | 0.5654 $\pm$ 0.0049 | 0.3569 $\pm$ 0.0029 |
| | Gated fusion | 0.5600 $\pm$ 0.0191 | 0.3640 $\pm$ 0.0104 |

Table 9: Comparison of RiNALMo-only and Graph-only (MLP) models on the mouse blind benchmark.

| RNA Type | Compartment | AUROC | | AUPRC | |
|---|---|---|---|---|---|
| | | RiNALMo-only | Graph-only (MLP) | RiNALMo-only | Graph-only (MLP) |
| **mRNA** | Nucleus | 0.6871 | 0.6573 | 0.6634 | 0.6429 |
| | Exosome | 0.4799 | 0.4970 | 0.1790 | 0.1906 |
| | Cytoplasm | 0.2972 | 0.3172 | 0.2523 | 0.2591 |
| | **Macro Avg** | **0.4881** | **0.4905** | **0.3649** | **0.3642** |
| **miRNA** | Nucleus | 0.5093 | 0.5580 | 0.2069 | 0.2356 |
| | Exosome | 0.5423 | 0.4456 | 0.8756 | 0.8211 |
| | Mitochondrion | 0.4838 | 0.5004 | 0.2944 | 0.2883 |
| | **Macro Avg** | **0.5118** | **0.5013** | **0.4589** | **0.4484** |
| **lncRNA** | Nucleus | 0.6626 | 0.5673 | 0.6312 | 0.5474 |
| | Exosome | 0.4669 | 0.4423 | 0.4726 | 0.4265 |
| | **Macro Avg** | **0.5648** | **0.5048** | **0.5519** | **0.4870** |

### Supplementary Material S3: RNA-Specific Processing Strategies

The time complexity for feature extraction is $O(n^2)$ for graph construction (where $n$ is sequence length), with optimizations for long sequences. RNA-specific adaptive processing preserves localization signals:

- **Short** ($\leq 1,000$ nt): Full processing - complete graph construction and feature extraction.
- **Medium** ($1,000 < L \leq 5,000$ nt): Smart truncation preserving 30% 5' region, 40% middle, 30% 3' region to retain UTRs and structural elements.
- **Long** ($> 5,000$ nt): Sliding window approach (window=500 nt, stride=250 nt) with weighted aggregation by motif density to capture distributed localization signals.

### References


Brandes, U. (2001). A faster algorithm for betweenness centrality. *Journal of Mathematical Sociology*, 25(2):163–177.

Chen, C.-Y. and Shyu, A.-B. (1995). Au-rich elements: characterization and importance in mrna degradation. *Trends in Biochemical Sciences*, 20(11):465–470.

Freier, S. M., Kierzek, R., Jaeger, J. A., Sugimoto, N., Caruthers, M. H., Neilson, T., and Turner, D. H. (1986). Improved free-energy parameters for predictions of rna duplex stability. *Proceedings of the National Academy of Sciences*, 83(24):9373–9377.

Hofacker, I. L., Fontana, W., Stadler, P. F., Bonhoeffer, L. S., Tacker, M., and Schuster, P. (1994). Fast folding and comparison of rna secondary structures. *Monatshefte für Chemie*, 125(2):167–188.

Lundberg, S. M. and Lee, S.-I. (2017). A unified approach to interpreting model predictions. In *Advances in Neural Information Processing Systems*, volume 30, pages 4765–4774.

Mathews, D. H., Sabina, J., Zuker, M., and Turner, D. H. (1999). Expanded sequence dependence of thermodynamic parameters improves prediction of rna secondary structure. *Journal of Molecular Biology*, 288(5):911–940.

Schneider, T. D. and Stephens, R. M. (1990). Sequence logos: a new way to display consensus sequences. *Nucleic Acids Research*, 18(20):6097–6100.

Turner, D. H. and Mathews, D. H. (2009). Nndb: the nearest neighbor parameter database for predicting stability of nucleic acid secondary structure. *Nucleic Acids Research*, 38(Database issue):D280–D282.

Valadi, H., Ekström, K., Bossios, A., Sjöstrand, M., Lee, J. J., and Lötvall, J. O. (2007). Exosome-mediated transfer of mrnas and micrornas is a novel mechanism of genetic exchange between cells. *Nature Cell Biology*, 9(6):654–659.

Yakovchuk, P., Protozanova, E., and Frank-Kamenetskii, M. D. (2006). Base-stacking and base-pairing contributions into thermal stability of the dna double helix. *Nucleic Acids Research*, 34(2):564–574.

Yan, Z., Hsu, C.-C., and Hsu, P.-W. (2016). A comprehensive analysis of rna sequences reveals the existence of a hidden layer of structural organization. *Scientific Reports*, 6:23129.

Zhang, B., Gunawardane, L., Niazi, F., Jahanbani, F., Chen, X., and Valadkhan, S. (2020). A novel rna motif mediates nuclear retention of a subset of long non-coding rnas. *Nucleic Acids Research*, 48(9):4812–4827.
